# A global synthesis reveals biodiversity-mediated benefits for crop production

**DOI:** 10.1101/554170

**Authors:** Matteo Dainese, Emily A. Martin, Marcelo A. Aizen, Matthias Albrecht, Ignasi Bartomeus, Riccardo Bommarco, Luisa G. Carvalheiro, Rebecca Chaplin-Kramer, Vesna Gagic, Lucas A. Garibaldi, Jaboury Ghazoul, Heather Grab, Mattias Jonsson, Daniel S. Karp, Christina M. Kennedy, David Kleijn, Claire Kremen, Douglas A. Landis, Deborah K. Letourneau, Lorenzo Marini, Katja Poveda, Romina Rader, Henrik G. Smith, Teja Tscharntke, Georg K.S. Andersson, Isabelle Badenhausser, Svenja Baensch, Antonio Diego M. Bezerra, Felix J.J.A. Bianchi, Virginie Boreux, Vincent Bretagnolle, Berta Caballero-Lopez, Pablo Cavigliasso, Aleksandar Ćetković, Natacha P. Chacoff, Alice Classen, Sarah Cusser, Felipe D. da Silva e Silva, G. Arjen de Groot, Jan H. Dudenhöffer, Johan Ekroos, Thijs Fijen, Pierre Franck, Breno M. Freitas, Michael P.D. Garratt, Claudio Gratton, Juliana Hipólito, Andrea Holzschuh, Lauren Hunt, Aaron L. Iverson, Shalene Jha, Tamar Keasar, Tania N. Kim, Miriam Kishinevsky, Björn K. Klatt, Alexandra-Maria Klein, Kristin M. Krewenka, Smitha Krishnan, Ashley E. Larsen, Claire Lavigne, Heidi Liere, Bea Maas, Rachel E. Mallinger, Eliana Martinez Pachon, Alejandra Martínez-Salinas, Timothy D. Meehan, Matthew G.E. Mitchell, Gonzalo A.R. Molina, Maike Nesper, Lovisa Nilsson, Megan E. O’Rourke, Marcell K. Peters, Milan Plećaš, Simon G. Potts, Davi de L. Ramos, Jay A. Rosenheim, Maj Rundlöf, Adrien Rusch, Agustín Sáez, Jeroen Scheper, Matthias Schleuning, Julia Schmack, Amber R. Sciligo, Colleen Seymour, Dara A. Stanley, Rebecca Stewart, Jane C. Stout, Louis Sutter, Mayura B. Takada, Hisatomo Taki, Giovanni Tamburini, Matthias Tschumi, Blandina F. Viana, Catrin Westphal, Bryony K. Willcox, Stephen D. Wratten, Akira Yoshioka, Carlos Zaragoza-Trello, Wei Zhang, Yi Zou, Ingolf Steffan-Dewenter

**Affiliations:** Institute for Alpine Environment, Eurac Research, Viale Druso 1, 39100 Bolzano, Italy; Department of Animal Ecology and Tropical Biology, Biocenter, University of Würzburg, Am Hubland, 97074 Würzburg, Germany; Grupo de Ecología de la Polinización - INIBIOMA (Universidad Nacional del Comahue–CONICET), 8400 Bariloche, Rio Negro, Argentina; Agroecology and Environment, Agroscope, Reckenholzstrasse 191, 8046 Zurich, Switzerland; Estación Biológica de Doñana (EBD-CSIC), Integrative Ecology. E-41092 Sevilla, Spain; Swedish University of Agricultural Sciences, Department of Ecology, 750 07 Uppsala, Sweden; Departamento de Ecologia, Universidade Federal de Goias (UFG), Goiânia, Brazil; Faculdade de Ciencias, Centre for Ecology, Evolution and Environmental Changes (CE3C), Universidade de Lisboa, Lisboa, Portugal; Natural Capital Project, Stanford University, Stanford, California, USA; CSIRO, GPO Box 2583, Brisbane, QLD 4001, Australia; Instituto de Investigaciones en Recursos Naturales, Agroecología y Desarrollo Rural (IRNAD), Sede Andina, Universidad Nacional de Río Negro (UNRN) y CONICET. Mitre 630, CP 8400, San Carlos de Bariloche, Río Negro, Argentina; Department of Environmental Systems Science, ETH Zurich, Universitätstrasse 16, 8092 Zurich, Switzerland; Cornell University, Department of Entomology, Ithaca, NY 14853, USA; Department of Wildlife, Fish, and Conservation Biology; University of California Davis, Davis, CA 95616, USA; Global Lands Program, The Nature Conservancy, 117 E. Mountain Avenue, Fort Collins, CO 80524, USA; Plant Ecology and Nature Conservation Group, Wageningen University, Droevendaalsesteeg 3a, Wageningen 6708 PB, The Netherlands; IRES and Biodiversity Research Centre, University of British Columbia, Vancouver, British Columbia, Canada; Michigan State University, Department of Entomology and Great Lakes Bioenergy Research Center, 204 CIPS, 578 Wilson Ave, East Lansing, MI 48824, USA; Department of Environmental Studies, University of California, Santa Cruz, CA USA 95064; DAFNAE, University of Padova, viale dell’Università 16, 35020, Legnaro, Padova, Italy; School of Environment and Rural Science, University of New England, Armidale, 2350, Australia; Centre for Environmental and Climate Research, Lund University, S-223 62 Lund, Sweden; Department of Biology, Lund University, S-223 62 Lund, Sweden; Agroecology, Department of Crop Sciences, University of Göttingen, Germany; USC1339 INRA-CNRS, CEBC UMR 7372, CNRS & Université de La Rochelle, Beauvoir sur Niort, 79360, France; INRA, URP3F Unité de Recherche Pluridisciplinaire Prairie et Plantes Fourragères, Lusignan, 86600, France; Functional Agrobiodiversity, Department of Crop Sciences, University of Göttingen, Germany; Departamento de Zootecnia – CCA, Universidade Federal do Ceará, 60.356-000, Fortaleza, CE, Brazil; Farming Systems Ecology, Wageningen University & Research, P.O. Box 430, 6700 AK Wageningen, The Netherlands; LTSER Zone Atelier Plaine & Val de Sevre, CEBC UMR 7372, CNRS & Université de La Rochelle, Beauvoir sur Niort, 79360, France; Arthropods Department, Natural Sciences Museum of Barcelona, 08003 Barcelona, Spain; Instituto Nacional de Tecnología Agropecuaria, Estación Experimental Concordia. Estacion Yuqueri y vias del Ferrocarril s/n (3200), Entre Rios. Argentina; Faculty of Biology, University of Belgrade, Studentski trg 16, 11000 Belgrade, Serbia; Instituto de Ecología Regional - IER (Universidad Nacional de Tucumán-CONICET), 4107 Yerba Buena, Tucumán, Argentina; W.K. Kellogg Biological Station, Michigan State University, Michigan, USA; Federal Institute of Education, Science and Technology of Mato Grosso, Campus of Barra do Garças/MT, 78600-000, Brazil; Center of Sustainable Development, University of Brasília (UnB) – Campus Darcy Ribeiro, Asa Norte, Brasília-DF, 70910-900, Brazil; Wageningen Environmental Research, Wageningen University & Research, P.O. Box 47, 6700 AA Wageningen, The Netherlands; Natural Resources Institute, University of Greenwich, Central Avenue, Chatham Maritime, Kent, ME44TB, UK; INRA, UR 1115, Plantes et Systèmes de culture horticoles, 84000 Avignon, France; Centre for Agri-Environmental Research, School of Agriculture, Policy and Development, Reading University, RG6 6AR, UK; Department of Entomology, University of Wisconsin, Madison, WI 53705, USA; Human-Environment Systems, Ecology, Evolution, and Behavior, Department of Biological Sciences, Boise State University, Boise, ID, USA; Department of Integrative Biology, University of Texas at Austin, 205 W 24th Street, 401 Biological Laboratories, Austin, TX 78712, USA; Department of Biology and Environment, University of Haifa - Oranim, TIvon 36006, Israel; Kansas State University Department of Entomology, 125 Waters Hall, Manhattan, KS 66503 USA; Department of Evolutionary and Environmental Biology, University of Haifa, 3498838 Haifa, Israel; Chair of Nature Conservation and Landscape Ecology, University of Freiburg, Tennenbacher Straße 4, 79106 Freiburg, Germany; Institute for Plant Science and Microbiology, University of Hamburg, Hamburg, Germany; Ashoka Trust for Research in Ecology and the Environment (ATREE), Bangalore, India; Bren School of Environmental Science & Management, University of California, Santa Barbara, CA 93106-5131, USA; Seattle University, Department of Environmental Studies, 901 12th Avenue, Seattle, WA 9812, USA; Department of Botany and Biodiversity Research, Division of Conservation Biology, Vegetation Ecology and Landscape Ecology, University of Vienna, Rennweg 14, 1030 Vienna, Austria; University of Florida, Department of Entomology and Nematology, 1881 Natural Area Drive, Gainesville, FL, USA 32601; Agrosavia, Centro de Investigación Obonuco. Km 5 vía Obonuco - Pasto, Nariño. Colombia; Agriculture, Livestock and Agroforestry Program, Tropical Agricultural Research and Higher Education Center (CATIE), Cartago, Turrialba, 30501, Costa Rica; National Audubon Society, Boulder, CO 80305, USA; Institute for Resources, Environment and Sustainability, University of British Columbia, Vancouver, British Columbia, Canada; Cátedra de Avicultura, Cunicultura y Apicultura - Facultad de Agronomía, Universidad de Buenos Aires - CABA (C1417DSE), Argentina; School of Plant and Environmental Sciences, Virginia Tech, Blacksburg, VA, USA; Department of Ecology, University of Brasília (UnB) - Campus Universitário Darcy Ribeiro, Brasília - DF, 70910- 900, Brazil; Department of Entomology and Nematology, University of California, Davis, CA 95616, USA; INRA, UMR 1065 Santé et Agroécologie du Vignoble, ISVV, Université de Bordeaux, Bordeaux Sciences Agro, F-33883 Villenave d’Ornon Cedex, France; INIBIOMA, Universidad Nacional del Comahue, CONICET, Quintral 1250, Bariloche (8400), Argentina; Senckenberg Biodiversity and Climate Research Centre (SBiK-F), Senckenberganlage 25, 60325 Frankfurt am Main, Germany; Centre for Biodiversity and Biosecurity, University of Auckland, Auckland, New Zealand; Department of Environmental Science, Policy & Management, University of California Berkeley, USA; South African National Biodiversity Institute, Kirstenbosch Research Centre, Private Bag X7, Claremont, 7735, South Africa; School of Agriculture and Food Science and Earth Institute, University College Dublin, Belfield, Dublin 4, Ireland; School of Natural Sciences, Trinity College Dublin, Dublin 2, Ireland; Institute for Sustainable Agro-Ecosystem Services, School of Agriculture and Life Sciences, The University of Tokyo, 188-0002 Tokyo, Japan; Forestry and Forest Products Research Institute, 1 Matsunosato, Tsukuba, Ibaraki 305-8687, Japan; Instituto de Biologia, Universidade Federal da Bahia, 40170-210 Salvador, Brazil; Bio-Protection Research Centre, Lincoln University, Lincoln, New Zealand; Fukushima Branch, National Institute for Environmental Studies, Japan; Environment and Production Technology Division, International food policy research institute, Washington DC, USA; Department of Health and Environmental Sciences, Xi’an-Jiaotong Liverpool University, Suzhou, China

## Abstract

Human land use threatens global biodiversity and compromises multiple ecosystem functions critical to food production. Whether crop yield-related ecosystem services can be maintained by few abundant species or rely on high richness remains unclear. Using a global database from 89 crop systems, we partition the relative importance of abundance and species richness for pollination, biological pest control and final yields in the context of on-going land-use change. Pollinator and enemy richness directly supported ecosystem services independent of abundance. Up to 50% of the negative effects of landscape simplification on ecosystem services was due to richness losses of service-providing organisms, with negative consequences for crop yields. Maintaining the biodiversity of ecosystem service providers is therefore vital to sustain the flow of key agroecosystem benefits to society.

## INTRODUCTION

Natural and modified ecosystems contribute a multitude of functions and services that support human well-being (*1, 2*). It has long been recognized that biodiversity plays an important role in the functioning of ecosystems (*3*), but the dependence of ecosystem services on biodiversity is under debate. An early synthesis revealed inconsistent results (*4*), whereas subsequent studies suggest that a few dominant species may supply the majority of ecosystem services (*5, 6*). It thus remains unclear whether a few dominant or many complementary species are needed to supply ecosystem services. A major limitation to resolving these relationships is a lack of evidence from real-world human-driven biodiversity changes (*7, 8*). For instance, changes in richness or abundance of service-providing organisms in response to land-clearing for agriculture (*9, 10*), could alter the flow of benefits to people.

Over the past half-century, the need to feed a growing world population has led to dramatically expanded and intensified agricultural production, transforming many regions into simplified landscapes (*11*). This transformation has contributed to enhanced agricultural production, but has also led to the degradation of the global environment. The loss of biodiversity can disrupt key intermediate services to agriculture, such as crop pollination (*12*) and biological pest control (*13*), that underpin the final provisioning service of crop production (*14*). Indeed, the recent stagnation or even decline of crop yields with ongoing intensification (*15*) indicates which alternative pathways are necessary to maintain future stable and sustainable crop production (*16*–*18*). An improved understanding of global biodiversity-driven ecosystem services in agroecosystems and their cascading effects on crop production, is urgently needed to forecast future supplies of ecosystem services and to pursue strategies for sustainable management (*8*).

We compiled an extensive database comprising 89 crop systems that measured richness and abundance of pollinators and pest natural enemies, and associated ecosystem services at 1,475 sampling locations around the world (Fig. 1A). Our study is focused on the ecosystem services of pollination and biological pest control, because these services are essential to crop production and have been the focus of much research in recent decades (*1*). We quantified pollinator and pest natural enemy richness as the number of unique taxa sampled from each location (field), and abundance as the sum of individuals sampled per field. We calculated a standardized index of pollination services using measures of pollination success and plant reproduction, and of pest control services using measures of natural enemy activity and crop damage (*19*). We also characterized the 1-km landscape surrounding each field by measuring the percentage of cropland and used this metric as a measure of landscape simplification (*20, 21*). Using a Bayesian multilevel modelling approach, we addressed three fundamental, yet unresolved questions in the biodiversity-ecosystem function framework: (i) are pollinator and natural enemy richness consistently related to pollination and pest control services independent of abundance?; (ii) does landscape simplification indirectly impact ecosystem services mediated by a loss of local community diversity?; and lastly, (iii) how strong are the cascading effects of landscape simplification on final crop production?

**Fig. 1.**
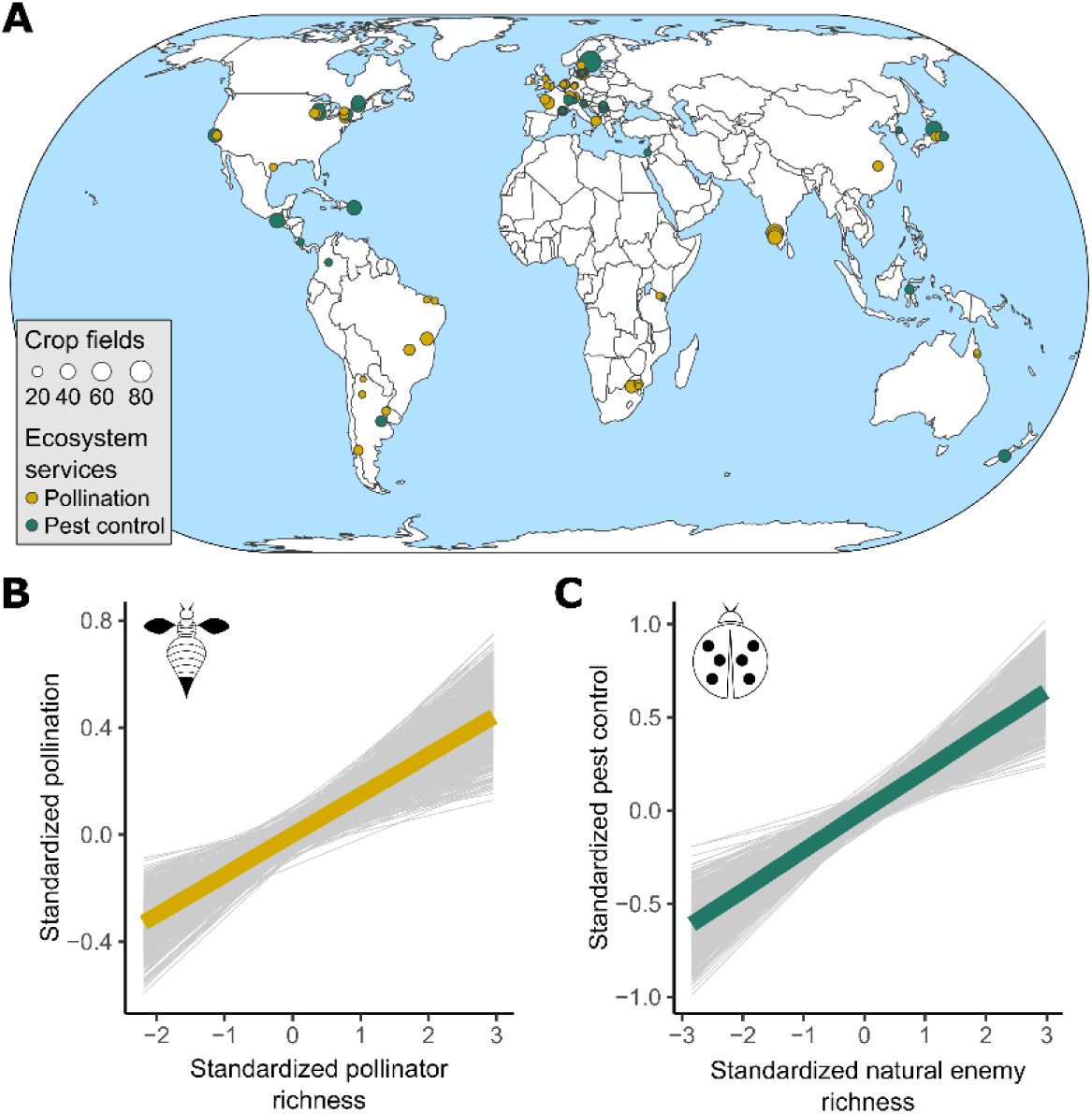
Distribution of analyzed crop systems and effects of richness on ecosystem services provisioning. (**A**) Global distribution of the 89 crop systems. Crop systems were defined as a given crop species, in a particular region and year (further details of crop systems are given in table S1). (**B**) Global effect of pollinator richness on pollination (*N* = 821 fields of 52 crop systems). (**C**) Global effect of natural enemy richness on pest control (*N* = 654 fields of 37 crop systems). The thick line in each plot represents the median of the posterior distribution of the model. Light grey lines represent 1,000 random draws from the posterior. The lines are included to depict uncertainty of the modelled relationship.

## RESULT AND DISCUSSION

Importantly, we found clear evidence that richness of service-providing organisms positively influenced ecosystem service delivery. This was detected for both pollination (Fig. 1B and table S2) and pest control (Fig. 1C and table S2), and in almost all crop systems (figs. S1 and S2). By a path analysis – where we tested the assumption that richness drives abundance resource use in addition to the classic view where richness is a function of the local community size (*19, 22*) – we further showed that these positive relationships were determined by both direct and mediated effects of richness and abundance of service-providing organisms on ecosystem services (fig. S3, table S3 and S4). The integration of different aspects of community structure in a single analysis revealed a more multilayered relationship between biodiversity and ecosystem services than has been previously acknowledged. These results complement previous findings for pollination (*6, 14, 23*) and pest control (*24*) and indicate that: (i) both richness and abundance contribute to support these two key ecosystem services in agriculture; and (ii) abundance and richness influence each other and cannot be interpreted in isolation. Hence, we find strong support for the role of species-rich communities in supporting pollination and pest control services.

Further, we found that landscape simplification indirectly affected ecosystem services by reducing the richness of service-providing organisms. Roughly a third of the negative effects of landscape simplification on pollination were due to a loss in pollinator richness (Fig. 2A and table S5). This effect was even greater for pest control where natural enemy richness mediated about 50% of the total effect of landscape simplification (Fig. 2B and table S5). A consistent richness-mediated effect was also confirmed when we tested the direct and indirect effects of landscape simplification on ecosystem services via changes in both richness and abundance (fig. S4 and table S6). Importantly, the effect of landscape simplification on ecosystem services was minimized when not considering the mediated effect of richness, especially for pest control. Indeed, we did not find a direct landscape simplification effect on pest control (all highest density intervals overlapped zero; Fig. 2B and table S5). Together, these results demonstrate strong negative indirect effects of landscape simplification on biodiversity-driven ecosystem services in agroecosystems, and the importance of the richness of service-providing organisms in mediating these effects.

**Fig. 2.**
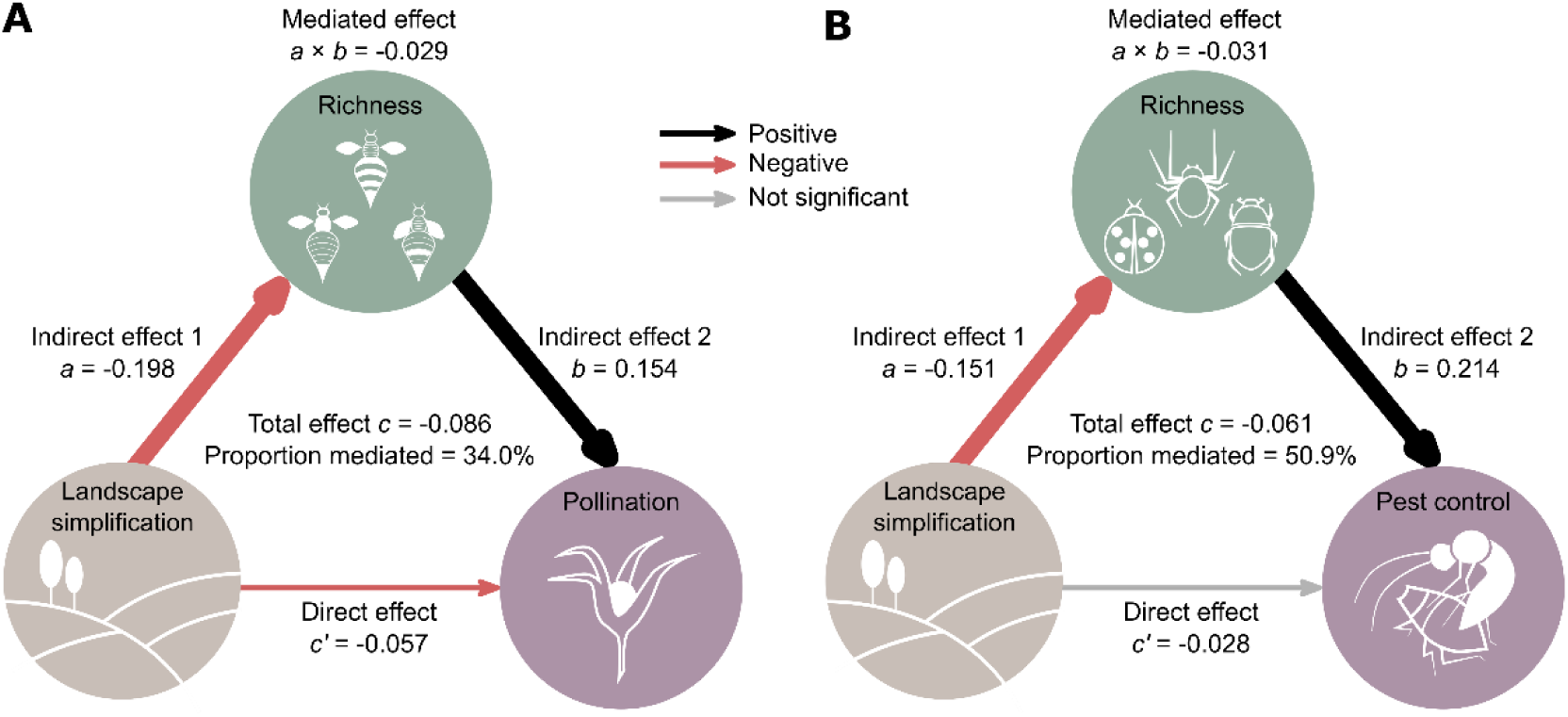
Direct and indirect effects of landscape simplification on richness of service-providing organisms and associated ecosystem services. (**A**) Path model of landscape simplification as a predictor of pollination, mediated by pollinator richness (*N* = 821 fields of 52 crop systems). (**B**) Path model of landscape simplification as a predictor of pest control, mediated by natural enemy richness (*N* = 654 fields of 37 crop systems). Coefficients of the three causal paths (*a, b, c*’) correspond to the median of the posterior distribution of the model. The proportion mediated is the mediated effect (*a* × *b*) divided by the total effect (*c*). Black and red arrows represent positive or negative effects, respectively. Arrow widths are proportional to highest density intervals (HDIs). Grey arrows represent non-significant effects (HDIs overlapped zero).

Finally, for a subset of the data that had crop production information (676 fields of 42 crop systems) we found that the cascading effects of landscape simplification mediated through richness and associated ecosystem services led to lower crop production. This was detected for both pollination (Fig. 3A and table S7) and pest control (Fig. 3B and table S7). Specifically, landscape simplification reduced both pollinator and natural enemy richness which had indirect consequences for pollination and pest control and, in turn, decreased crop production. Pollinator abundance was also negatively affected by landscape simplification, but in contrast to richness, abundance had no significant effect on pollination services (Fig. 3A). Effects of landscape simplification on natural enemy abundance were even weaker (Fig. 3B). For pest control, a positive link with crop production was detected in fields where the study area was not sprayed with insecticides during the course of the experiment (Fig. 3B), but not when considering all sites combined (with and without insecticide use; fig. S5). In sprayed areas, we did not find a pest control effect (all highest density intervals overlapped zero), probably because effects were masked by insecticide use (*25, 26*). Importantly, a positive link with crop production was detected even though measures used to estimate pest control (natural enemy and pest activity) were not direct components of crop production, as was the case of pollination measures (fruit or seed set). Though only available from a subset of the data, this result underpins that the effects of landscape simplification can cascade up to reducing the final provisioning service of crop production.

**Fig. 3.**
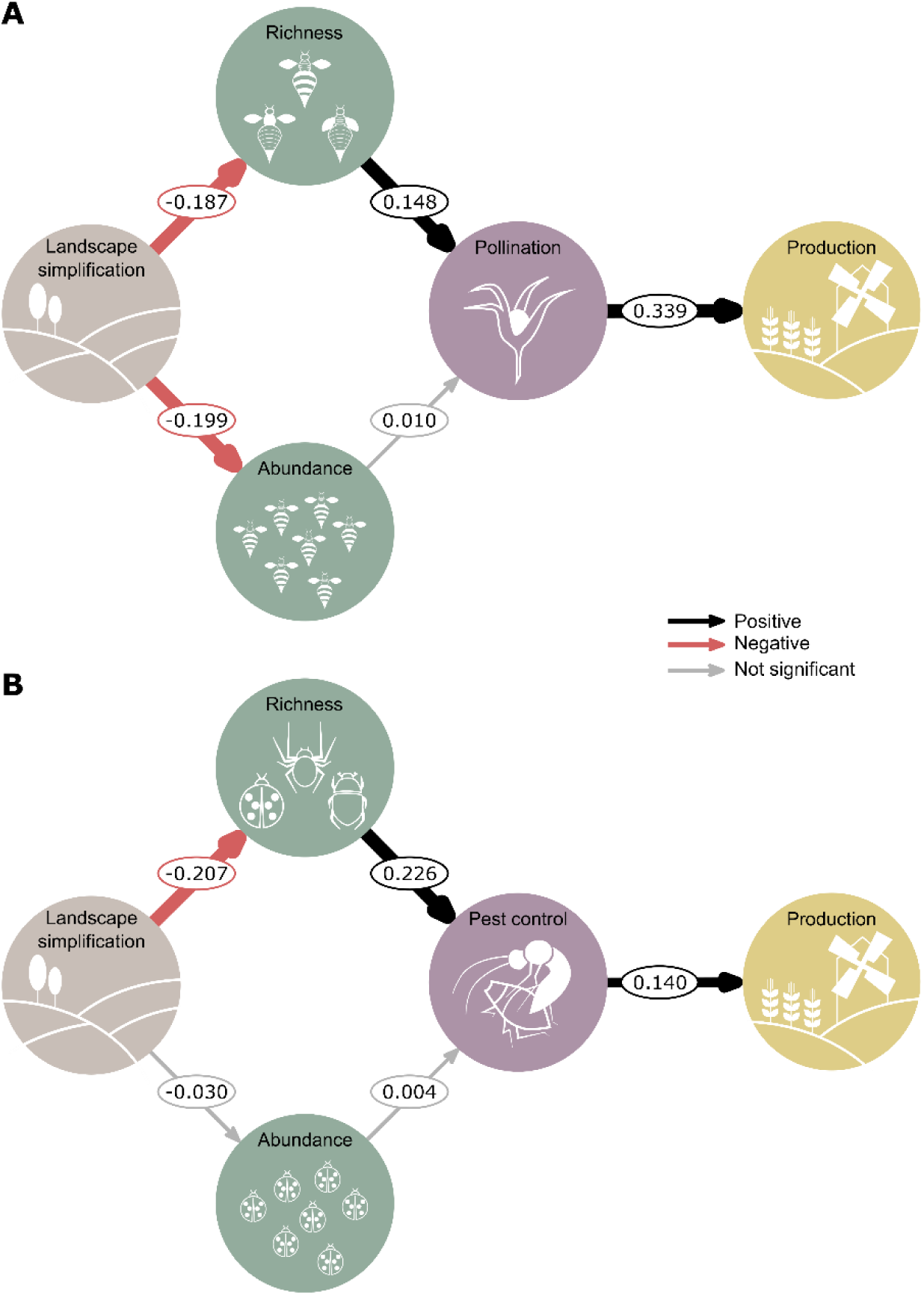
Direct and cascading effects of landscape simplification on final crop production via changes in richness, abundance and ecosystem services. (**A**) Path model representing direct and indirect effects of landscape simplification on final crop production through changes in pollinator richness, abundance and pollination (*N* = 440 fields of 27 crop systems). (**B**) Path model representing direct and indirect effects of landscape simplification on final crop production through changes in natural enemy richness, abundance and pest control (only insecticide-free areas were considered in the model (*N* = 184 fields of 14 crop systems). Path coefficients are effect sizes estimated from the median of the posterior distribution of the model. Black and red arrows represent positive or negative effects, respectively. Arrow widths are proportional to highest density intervals (HDIs). Grey arrows represent non-significant effects (HDIs overlapped zero).

Our findings suggest that some previously inconsistent responses of natural enemy abundance and activity to surrounding landscape composition (*27*) can be reconciled by considering richness in addition to abundance. Although richness and abundance are often correlated, their response to environmental variation can differ. This was evident in the path analysis showing a strong effect of landscape simplification on richness, but only a marginal effect on abundance (fig. S4b). Moreover, effect sizes for natural enemy abundances in individual crop systems (fig. S6) showed similarly inconsistent responses to the previous synthesis (*27*). For pest control, both results are instead well aligned (Fig. 2B).

Using an integrative model to assess key ecological theory, we demonstrate that the negative effects of landscape simplification on service supply and final crop production are primarily mediated by loss of species. We found strong evidence for positive biodiversity-ecosystem service relationships, highlighting that managing landscapes to enhance the richness of service-providing organisms (*28*) is a promising pathway towards a more sustainable food production globally. In an era of rapid environmental changes, preserving biodiversity-driven services will consistently confer greater resilience to agroecosystems, such that we could expect improved crop production under a broader range of potential future conditions.

## MATERIALS AND METHODS

### Database compilation

We compiled data from crop studies where measures of richness and abundance of service-providing organisms (pollinators or natural enemies) and associated ecosystem services (pollination and biological pest control) were available for the same sites. If available, we also included information on yield. Studies were identified by first searching the reference lists of recent meta-analyses (*6, 14, 27, 29, 30*) and then directly contacting researchers. For pest control, data were mostly provided from a recent pest control database (*27*). Of 191 researchers initially contacted, 86 provided data that met our criteria. Overall, we analyzed data from 89 crop systems and 1,475 fields in 27 countries around the world (table S1). Crop systems were defined as a given crop species, in a particular region and year (*14*). Twenty-nine crops were considered, including a wide array of annual and perennial fruit, seed, nut, stimulant, pulse, cereal and oilseed crops. Crop systems represented the spectrum of management practices, that is, conventional, low-input conventional, integrated pest management and organic farming. In 76% of fields, pest control experiments were performed in insecticide-free areas. In some fields this information was not available (7%) or insecticides were applied (17%). As similar studies were frequently performed in the same area, occasionally in the same year, and studies with multiple years usually used different sites each year, we did not nest year within study. Instead, we considered each year of multi-year studies (that is, 10 studies) to be an independent dataset and used study-year combinations as the highest hierarchical unit.

### Pollinator and pest natural enemy richness and abundance

Studies used a broad range of methods, which we categorized as active or passive (*31*) to sample pollinators or natural enemies. Active sampling methods included netting pollinators seen on crop flowers, hand-collecting individuals on plants, observational counting, sweep-netting, and vacuum sampling. Passive sampling methods were malaise traps, pan traps, pitfall traps, and sticky cards. Active sampling was performed in 85% of pollinator sampling fields and in 50% of natural enemy sampling fields.

Pollinators included representatives from the orders Hymenoptera, Diptera, Lepidoptera, and Coleoptera. Bees (Hymenoptera: Apoidea) were the most commonly observed pollinators and included *Apis* bees (Apidae: *Apis mellifera, Apis cerana, Apis dorsata, Apis florea*), stingless bees (Apidae: Meliponini), bumble bees (Apidae: *Bombus* spp.), carpenter bees (Apidae: Xylocopini), small carpenter bees (Apidae: Ceratinini), sweat bees (Halictidae), long-horned bees (Apidae: Eucerini), plasterer bees (Colletidae), mining bees (Andrenidae), and mason bees (Megachilidae). Non-bee taxa included syrphid flies (Diptera: Syrphidae), other flies (Diptera: Calliphoridae, Tachinidae, and Muscidae), butterflies and moths (Lepidoptera), various beetle families (Coleoptera) and hymenopterans including ants (Formicidae) and the paraphyletic group of non-bee aculeate wasps.

Natural enemies included ground beetles (Coleoptera), flies (Diptera), spiders (Aranea), hymenopterans including ants (Formicidae) and wasps, bugs (Hemiptera), thrips (Thysanoptera), net-winged insects (Neuroptera), bats and birds.

We calculated pollinator and natural enemy richness as the number of unique taxa sampled per crop system, method and field. A taxon was defined as a single biological type (that is, species, morphospecies, genus, family) determined at the finest taxonomic resolution to which each organism was identified. In almost 70% of cases, taxonomic resolution was to species-level (averaged proportion among all studies), but sometimes it was based on morphospecies- (15%), genus- (8%) or family-levels (7%). Taxon richness per field varied between 1 and 49 for pollinators and between 1 and 40 for natural enemies. Abundance reflects the sum of individuals sampled per crop system, method and field. Pollinator richness and abundance were calculated either including or excluding honey bees (*Apis mellifera*). *Apis mellifera* was considered as the only species within the honey bee group for consistency across all datasets (*30*). Other *Apis* bees (that is, *Apis cerana, Apis dorsata, Apis florea*) were not pooled into the honey bee category as the large majority of observed individuals are derived from feral populations. Feral and managed honey bees were analysed as a single group because they cannot be distinguished during field observations. Feral honey bees were uncommon in most studies except for those in Africa and South America (*Apis mellifera* is native in Africa, while it was introduced to the Americas). In studies with subsamples within a field (that is, plots within fields or multiple sampling rounds within fields), we calculated the total number of individuals and unique taxa across these subsamples.

### Pollination and pest control services

As different methods were used to quantify pollination or pest control services across studies, standardization was necessary to put all the indices on equivalent terms. Therefore, we transformed each index *y* in each field *i* in each crop system *j* using *z*-scores. We preferred the used of *z*-scores over other transformations (for example, division by the maximum), because *z*-scores do not constrain the variability found in the raw data, as do other indices that are bounded between 0 and 1. We used the proportion of flowers that set fruit (that is, fruit set), the average number of seeds per fruits (that is, seed set), or the estimated measures of pollinator contribution to plant reproduction (that is, differences in fruit weight between plants with and without insect pollination, hereafter Δ fruit weight) as measures of pollination services. We then converted these measures into the pollination index. The pest control index was calculated using measures of natural enemy activity or pest activity. Natural enemy activity was measured by sentinel pest experiments where pests were placed in crop fields and predation or parasitism rates were monitored, or field exclosure experiments where cages were used to exclude natural enemies to quantify differences in pest abundance or crop damage between plants with and without natural enemies. Pest activity was measured as the fraction or amount of each crop consumed, infested, or damaged. We inverted standardized values of pest activity by multiplying by −1, as low values indicate positive contributions to the ecosystem service.

### Crop production

Depending on the crop type, marketable crop yield is not only valued by farmers in terms of area-based yield, but also in terms of fruit or seed weight [for example, in coffee, sunflower or strawberry fields; (*32, 33*)] or seed production per plant [for example, in seed production fields; (*34*)]. Moreover, area-based yield and within-plant yield are often correlated (*35, 36*). Thus, we used both area-based yield and within-plant yield as measures of final crop production. Within-plant yield was measured by the total number (or mass) of seeds or fruits per plant, or by fruit or seed weight. Also in this case, we standardized variables (*z*-scores) to put all the indices on equivalent terms.

### Landscape simplification

Landscapes were characterized by calculating the percentage of cropland (annual and perennial) within a 1 km radius around the center of each crop field. This landscape metric has been used as a relevant proxy for characterizing landscape simplification (*20, 21*) and is often correlated with other indicators of landscape complexity (*37, 38*). Moreover, we used this metric because cropland data are readily accessible from publicly available land cover data and are more accurate than other land use types such as forests and grasslands (*39*), especially when detailed maps are not available. The 1 km spatial extent was chosen to reflect the typical flight and foraging distances of many insects including pollinators (*40, 41*) and natural enemies (*42, 43*). For studies where this information was not supplied by the authors, land uses were digitized using GlobeLand30 (*44*), a high-resolution map of Earth’s land cover. The derived land-cover maps were verified and, if necessary, corrected using a visual inspection of satellite images (Google Earth®). We then calculated the percentage of cropland within the radius using Quantum GIS 2.18 (Open Source Geospatial Foundation Project, http://qgis.osgeo.org). The average percentage of cropland was 67.5% for pollination studies and 41.5% for natural enemy studies.

### Data analysis

#### Data standardization

Before performing the analyses, we standardized the predictors (abundance, richness and landscape simplification) using z-scores within each crop system. This standardization was necessary to allow comparisons between studies with differences in methodology and landscape ranges (*45*). By doing this the focus of our analysis is on within-study effects rather than between-study effects (*46, 47*).

#### Relationship between richness and ecosystem services

The relationship between richness of service-providing organisms and related ecosystem services (Fig. 1B and 1C) was estimated from a Bayesian multilevel (partial pooling) model that allowed the intercept and the slope to vary among crop systems (also commonly referred to as random intercepts and slopes), following the equation:

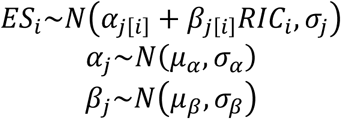

where *ES*_***i***_ is the ecosystem service index (pollination or pest control depending on the model), *RIC*_*i*_ is richness of service-providing organisms (pollinator or natural enemy richness depending on the model), and *j*_[*i*]_ represents observation *i* of crop system *j*. This partial-pooling model estimates both crop system-level responses [yielding an estimate for each crop system (*β*_*j*_)] and the distribution from which the crop system-level estimates are drawn, yielding a higher-level estimate of the overall response across crop systems (*μ*_*β*_). In addition, it accounts for variation in variance and sample size across observations (for example, crop systems, studies). The intercepts *α*_*j*_ and slopes *β*_*j*_ varied between crop systems according to a normal distribution with mean *μ* and standard deviation *σ*. Independent within-crop system errors also followed a normal distribution *ε*_*i*_ ∼ *N*(0,*σ*). We used weakly informative priors: Normal (0,10) for the population-level parameters (*α, β*) and half-Student-*t* (3, 0, 5) for the group-level standard deviation and residual standard deviation.

#### Direct and mediated effects of richness and abundance on service provisioning

As natural communities vary not only in number of species but also in number of individuals (abundance), it is important to incorporate these attributes when assessing or modelling biodiversity effects (*48, 49*). According to a revised version of the ‘more individuals hypothesis’ (*22*), we cannot necessarily infer that an increase in the number of individuals of a community causes an increase in the number of species in a unidirectional way, but theory also indicates that more species can exploit more diverse resources and may therefore maintain more individuals than species-poor communities. In a Bayesian multivariate response model with causal mediation effects (hereafter, mediation model), a form of path analysis, we thus verified two alternative paths between richness, abundance and ecosystem services. We tested (i) whether richness *per se* directly influences ecosystem services or is instead mediated by abundance (fig. S3a, b), and (ii) whether abundance *per se* directly influences ecosystem services or is instead mediated by richness (fig. S3c, d). Prior to analysis, we checked for data collinearity among abundance and richness by calculating the variance inflation factor (VIF). No signal of collinearity was detected in either model (VIFs were below 1.5). Mediation analysis is a statistical procedure to test whether the effect of an independent variable X on a dependent variable Y (X → Y) is at least partly explained via the inclusion of a third hypothetical variable, the mediator variable M (X → M → Y) (*50*). The three causal paths *a, b*, and *c*’ correspond to X’s effect on M, M’s effect on Y, and X’s effect on Y having taken M into account, respectively. The three causal paths correspond to parameters from two regression models, one in which M is the outcome and X the predictor, and one in which Y is the outcome and X and M the simultaneous predictors (fig. S7). From these parameters, we can compute the mediation effect (the product ab; also known as the indirect effect), and the total effect of X on Y,

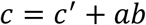

Thus, the total causal effect of X, which is captured by the parameter *c*, can be decomposed precisely into two components, a direct effect *c*’ and an indirect (mediation) effect *ab* (the product of paths *a* and *b*). To illustrate we first show the univariate multilevel (partial pooling) models following these equations:

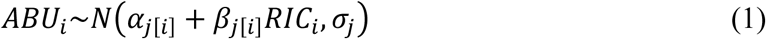

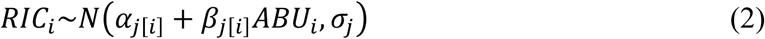

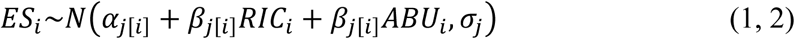

where is *ABU*_*i*_ abundance, *RIC*_*i*_ is richness of service-providing organisms, *ES*_*i*_ is the ecosystem service index, and the index *j*_[*i*]_ represents observation *i* of crop system *j*. We specified both multivariate multilevel models in a matrix-vector notion (*45*), as follows:

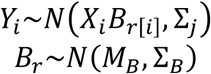

where *Y*_*i*_ is the matrix of response variables with observations *i* as rows and variables *r* as columns, *X*_*i*_ is the matrix of all predictors for response *r, B*_*r*_ are the regression parameters (*α* and *β*) for response *r, M*_*B*_ represents the mean of the distribution of the regression parameters, and *Σ*_*B*_ is the covariance matrix representing the variation of the regression parameters in the population groups. We used weakly informative priors: Normal (0,10) for the population-level parameters (*α, β*) and half-Student-t (3, 0, 5) for the group-levels standard deviation and residual standard deviation. In building the model, we ensured that no residual correlation between *ES*_*i*_ and *ABU*_*i*_ or *ES*_*i*_ and *RIC*_*i*_ was estimated [see ‘*set_rescor*’ function in the package *brms*; (*51*)]. The mediation analysis was implemented using the R package *sjstats* [v 0.15.0; (*52*)].

#### Direct and indirect effects of landscape simplification on ecosystem services

To estimate the direct and indirect effects of landscape simplification on richness and associated ecosystem services, we employed two models. First, we developed a mediation model to test whether landscape simplification directly influences ecosystem services or is mediated by richness. The model included the ecosystem service index as response, landscape simplification as predictor, and richness as mediator (Fig. 2). The separate regression models that made up the Bayesian multivariate multilevel model followed these equations:

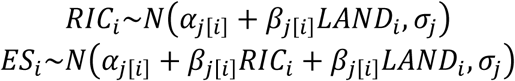

We then compiled a multilevel path analysis testing the direct and indirect effects of landscape simplification on ecosystem services via changes in both richness and abundance (fig. S4). The separate regression models that made up the model followed these equations:

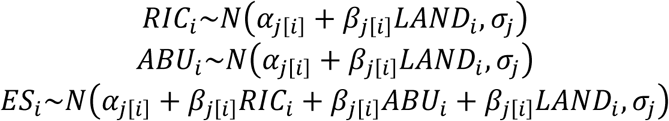

where *RIC*_*i*_ is richness of service-providing organisms, *LAND*_*i*_ is landscape simplification measured as the percentage of arable land surrounding each study site, *ABU*_*i*_ is abundance, *ES*_*i*_ is the ecosystem service index, and the index *j*_[*i*]_ represents observation *i* of crop system *j*. We then specified multivariate multilevel models in a matrix-vector notion, as explained above.

#### Cascading effects of landscape simplification on final crop production

For 42 crop systems and 676 fields (pollination model, *N* = 440 fields of 27 crop systems; pest control model, *N* = 236 fields of 15 crop systems; table S1), the data allowed us to employ a multilevel path analysis to examine cascading effects of landscape simplification on final crop production via changes in richness, abundance and ecosystem services. In this model, we expected that: (i) landscape simplification would have a direct effect on richness and abundance of service-providing organisms, (ii) richness and abundance of service-providing organisms would relate positively to intermediate services, which in turn, (iii) would increase final crop production (Fig. 3). The separate regression models that made up the path model followed these equations:

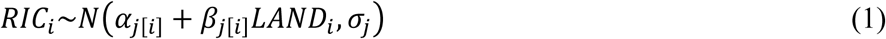

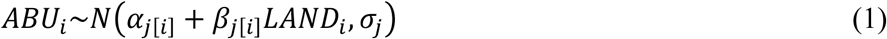

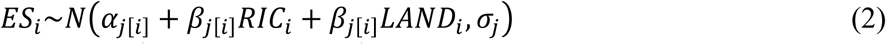

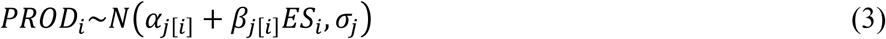

where *RIC*_*i*_ is richness of service-providing organisms, *LAND*_*i*_ is landscape simplification measured as the percentage of arable land surrounding each study site, *ABU*_*i*_ is abundance, *ES*_*i*_ is the ecosystem service index, *PROD*_*i*_ is crop production, the index *j*_[*i*]_ represents observation *i* of crop system *j*. We specified a multivariate multilevel model in a matrix-vector notion, as explained above.

#### Parameter estimation

All analyses were conducted in Stan through R (v. 3.4.3) using the package brms [v 2.2.0; (*51*)]. Stan provides efficient MCMC sampling via a No-U-Turn Hamiltonian Monte Carlo approach (*53*). Each model was run with four independent Markov chains of 5,000 iterations, discarding the first 2,500 iterations per chain as warm-up and resulting in 10,000 posterior samples overall. Convergence of the four chains and sufficient sampling of posterior distributions were confirmed by: (i) the visual inspection of parameter traces, (ii) ensuring a scale reduction factor 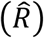 below 1.01, and (iii) effective size (*n*_*eff*_) of at least 10% of the number of iterations. For each model, posterior samples were summarized based on the Bayesian point estimate (median), standard error (median absolute deviation), and posterior uncertainty intervals by highest density intervals (HDIs), a type of credible interval which contains the required mass such that all points within the interval have a higher probability density than points outside the interval (*54*). The advantage of the Bayesian approach is the possibility not only to estimate expected values for each parameter, but also the uncertainty associated with these estimates (*55*). Thus, we calculated 80%, 90% and 95% HDIs for parameter estimates.

#### Sensitivity analyses

Given that different methods were used in different studies to quantify richness, ecosystem services and final crop production, we measured the sensitivity of our results to methodological differences.

(i) We verified whether treating each annual data set from multi-year studies separately could incorrectly account for the dependence of the data. We refitted the model testing the relationship between richness and ecosystem services including year nested within crop system (that is, crop system defined as crop-region combination). Then, we compared models (year-independent model vs. year-nested model) using leave-one-out cross-validation (LOO), a fully Bayesian model selection procedure for estimating pointwise out-of-sample prediction accuracy (*56*). We calculated the expected log pointwise predictive density 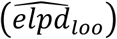, using the log-likelihood evaluated at the posterior simulations of the parameter values. Model comparison was implemented using R package loo [v 2.0.0; (*57*)]. We found that the year-nested model had a lower average predictive accuracy than the year-independent model for both pollination 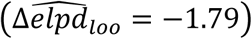 and pest control 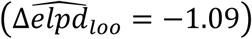, and therefore retained the year-independent model in our analysis.

(ii) We verified whether taxonomic resolution influenced the interpretation of results. We recalculated richness considering only organisms classified at the fine taxonomy level (species- or morphospecies-levels) and refitted the model testing the effect of richness on ecosystem services. We found no evidence that taxonomic resolution influenced our results. With a fine taxonomic resolution, the effects of richness on ecosystem services (*β*_pollinators_ = 0.1535, 90% HDIs = 0.0967 to 0.2141; *β*_enemies_ = 0.2262, 90% HDIs = 0.1420 to 0.3022; table S2) were nearly identical to the estimates presented in the main text (*β*_pollinators_ = 0.1532, 90% HDIs = 0.0892 to 0.2058; *β*_enemies_ = 0.2132, 90% HDIs = 0.1451 to 0.2810; table S2).

(iii) We verified whether the sampling methods used to collect pollinators (active vs. passive sampling techniques) influenced the relationship between pollinator richness and pollination using Bayesian hypothesis testing (*51*). Passive methods do not directly capture flower visitors and may introduce some bias (for example, they may underestimate flower visitors). However, our estimate was not influenced by sampling method (the one-sided 90% credibility interval overlapped zero; table S9). In accordance with this finding, the evidence ratio showed that the hypothesis tested (that is, estimates of studies with active sampling > estimates of studies with passive sampling) was only 0.78 times more likely than the alternative hypothesis.

(iv) We verified whether methodological differences in measuring pollination and pest control services influenced the relationship between richness and ecosystem services. Using Bayesian hypothesis testing, we tested whether the estimates differed among methods. The two-sided 95% credibility interval overlapped zero in all comparisons (estimates did not differ significantly; table S10) indicating that our estimate was not influenced by methodological differences in measuring ecosystem services. Furthermore, we tested effects including only inverted pest activity as a reflection of pest control. We found positive effects of natural enemy richness on inverted pest activity (*β* = 0.1307, 90% HDIs = 0.0102 to 0.2456), indicating that results were robust to the type of pest control measure considered.

(v) As honey bees are the most important and abundant flower visitors in some locations, we verified the potential influence of honey bees on our results by refitting the mediation model with honey bees. A positive direct contribution of richness to pollination was confirmed even after including honey bees (fig. S8). However, abundance was more important than richness when honey bees were considered.

(vi) Insecticide application during the course of the experiment could mask the effect of pest control on crop production (*25, 26*). We verified the potential influence of insecticide application on our results by refitting the model considering only fields where the study area was not sprayed with insecticide during the course of the experiment (*N* = 184 fields of 14 crop systems). Indeed, we found a pest control effect that was masked when considering all sites combined (with and without insecticide; fig. S5). We therefore show the insecticide-free model in the main text (Fig. 3).

(*vii*) We verified the consistency of our results considering only studies that measured area-based yield (sub-model). Only significant terms were retained in a simplified model. We found no evident differences between the sub-model (fig. S9) and the full model presented in the main text (Fig. 3).

## ACKNOWLEDGMENTS

This work was funded by EU-FP7 LIBERATION (311781) and Biodiversa-FACCE ECODEAL (PCIN-2014–048). For all further acknowledgements see the Supplementary Materials. We thank R. Carloni for producing icons in Figures 1 to 3.

## AUTHOR CONTRIBUTIONS

M.D., E.A.M. and I.S.D. conceived the study. M.D. performed statistical analyses and wrote the manuscript draft. The authors named from E.A.M. to T.T and I.S.D. discussed and revised earlier versions of manuscript. The authors named from G.K.S.A. to Y.Z. are listed alphabetically, as they contributed equally in gathering field data and providing several important corrections to subsequent manuscript drafts.

## DATA AND MATERIALS AVAILABILITY

The data and R script files used in this study are available from the corresponding authors.

