## Supplementary Materials for "A global synthesis reveals biodiversity-mediated benefits for crop production"

Contents:

**Supplementary Figures**


**Fig. S1.** Forest plot of the effect of pollinator richness on pollination for individual crop systems.

**Fig. S2.** Forest plot of the effect of natural enemy richness on pest control for individual crop systems.

**Fig. S3.** Direct and mediated effects of richness and abundance on ecosystem services.

**Fig. S4.** Direct and indirect landscape simplification effects on ecosystem services via changes in richness and abundance.

**Fig. S5.** Direct and cascading landscape simplification effects on final crop production via changes in natural enemy richness, abundance and pest control (all sites together, with and without insecticide application).

**Fig. S6.** Forest plot of the effect of landscape simplification on natural enemy abundance for individual crop systems.

**Fig. S7.** Mediation model.

**Fig. S8.** Direct and mediated effects of pollinator richness and abundance (with honey bees) on pollination.

**Fig. S9.** Direct and cascading landscape simplification effects on area-based yield via changes in richness, abundance and ecosystem services


**Supplementary Tables**


**Table S1.** List of 88 crop systems considered in our analyses.

**Table S2.** Model output for richness-ecosystem service relationships.

**Table S3.** Model output for path models testing direct and indirect effects (mediated by changes in abundance) of richness on ecosystem services.

**Table S4.** Model output for path models testing direct and indirect effects (mediated by changes in richness) of abundance on ecosystem services.

**Table S5.** Model output for path models testing direct and indirect effects (mediated by changes in richness) of landscape simplification on ecosystem services.

**Table S6.** Model output for path models testing the direct and cascading landscape simplification effects on ecosystem services via changes in richness and abundance.

**Table S7.** Model output for path models testing the direct and cascading landscape simplification effects on final crop production via changes in richness, abundance and ecosystem services.

**Table S8.** Model output for path models testing direct and mediated effects of pollinator richness and abundance (with honey bees) on pollination.

**Table S9.** Results of pairwise comparison of richness-ecosystem service relationships according to the methods used to sample pollinators and natural enemies.

**Table S10.** Results of pairwise comparison of richness-ecosystem service relationships according to the methods used to quantify pollination and pest control services.

**Supplementary Text**

Detailed Acknowledgemnts

**References**

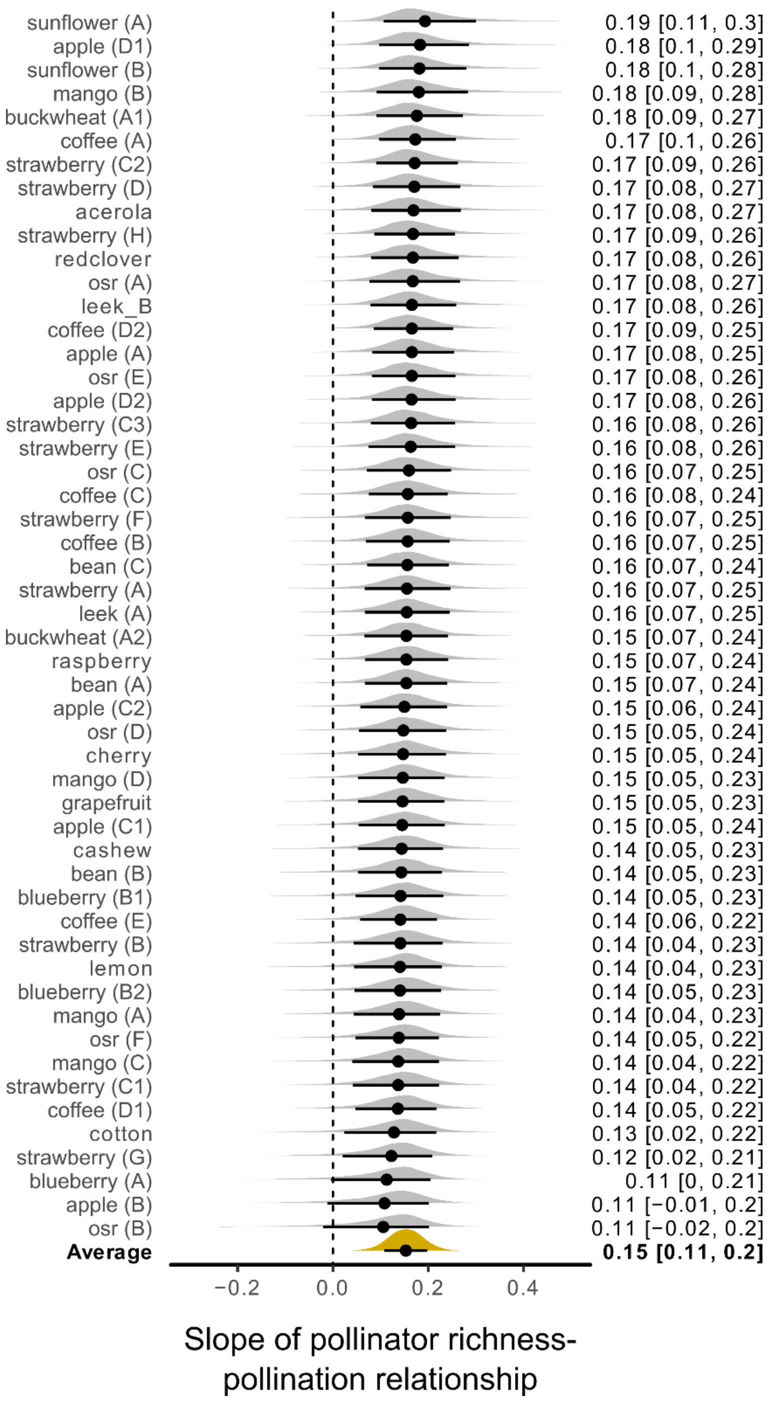


**Fig. S1.**

**Forest plot of the effect of pollinator richness on pollination for individual crop systems**. Each posterior distribution represents medians (symbol centres) and 90% density intervals (black lines).

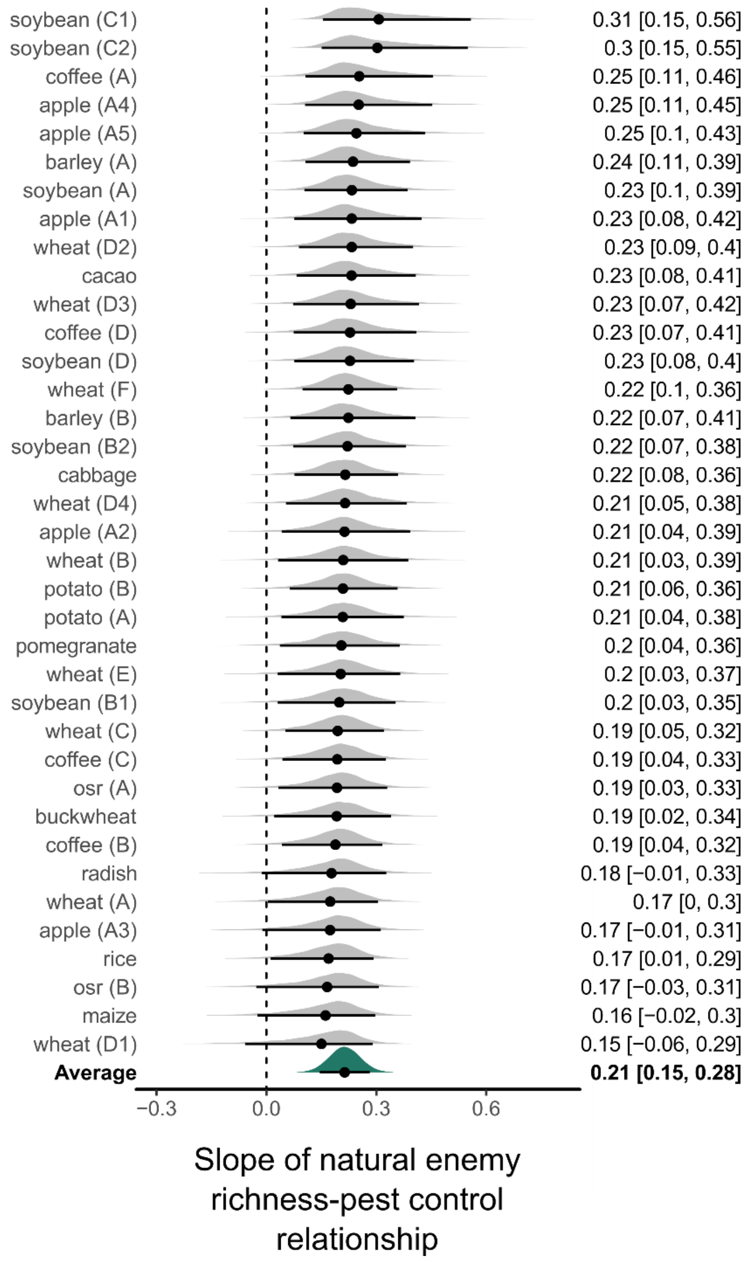


**Fig. S2.**

**Forest plot of the effect of natural enemy richness on pest control for individual crop systems.** Each posterior distribution represents medians (symbol centres) and 90% density intervals (black lines).

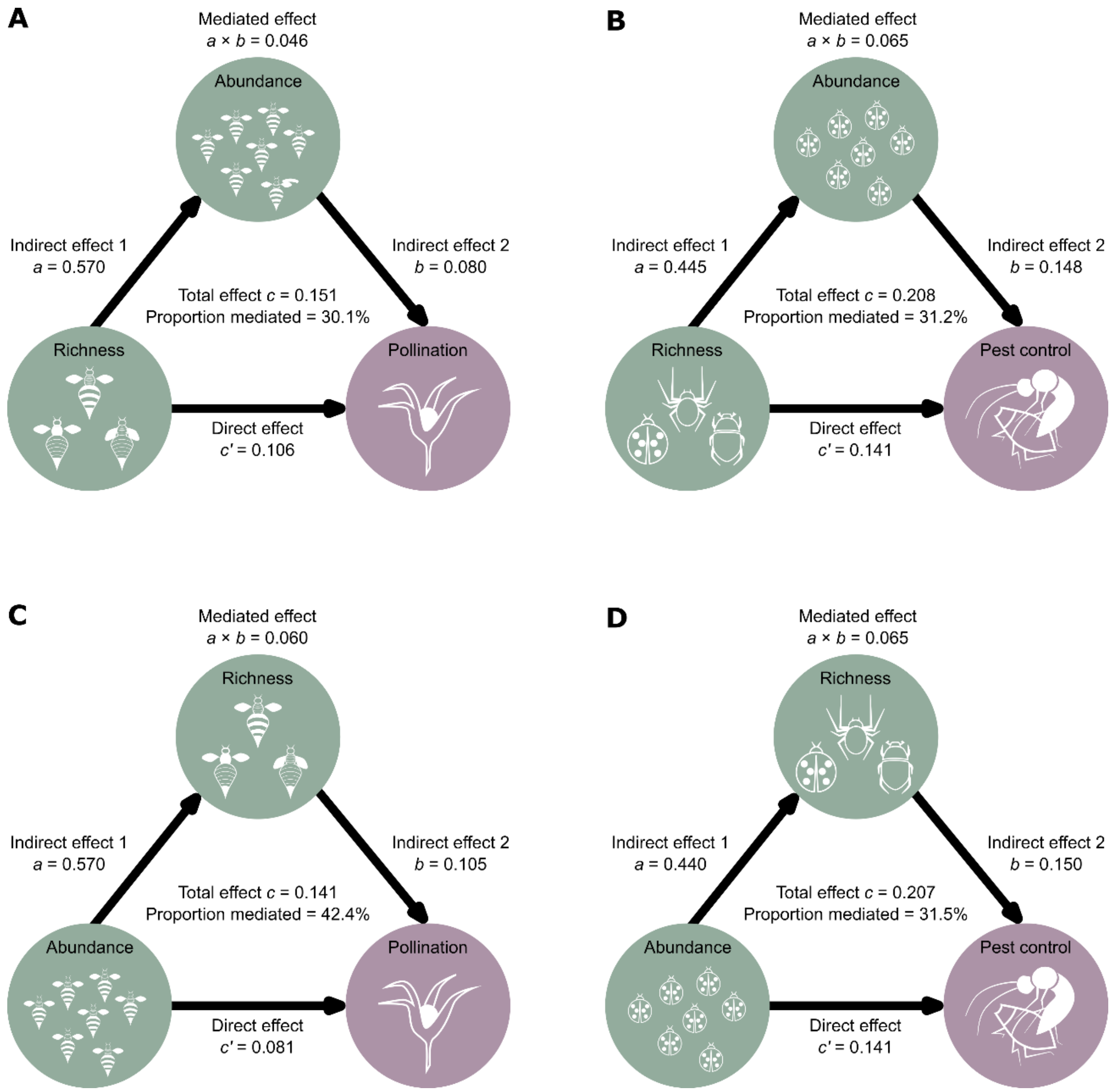


**Fig. S3.**

**Direct and mediated effects of richness and abundance on ecosystem services.** (**A**) Path model of pollinator richness as predictor of pollination, mediated by pollinator abundance. (**B**) Path model of natural enemy richness as predictor of pest control, mediated by natural enemy abundance. (**C**) Path model of pollinator abundance as predictor of pollination, mediated by pollinator richness. (**D**) Path model of natural enemy abundance as predictor of pest control, mediated by natural enemy richness. Pollination model, $N$ = 821 fields of 52 crop systems; pest control model, $N$ = 654 fields of 37 crop systems. Coefficients of the three causal paths ($a$, $b$, $c'$) correspond to the median of the posterior distribution of the model. The proportion mediated is the mediated effect ($a \times b$) divided by the total effect ($c$).

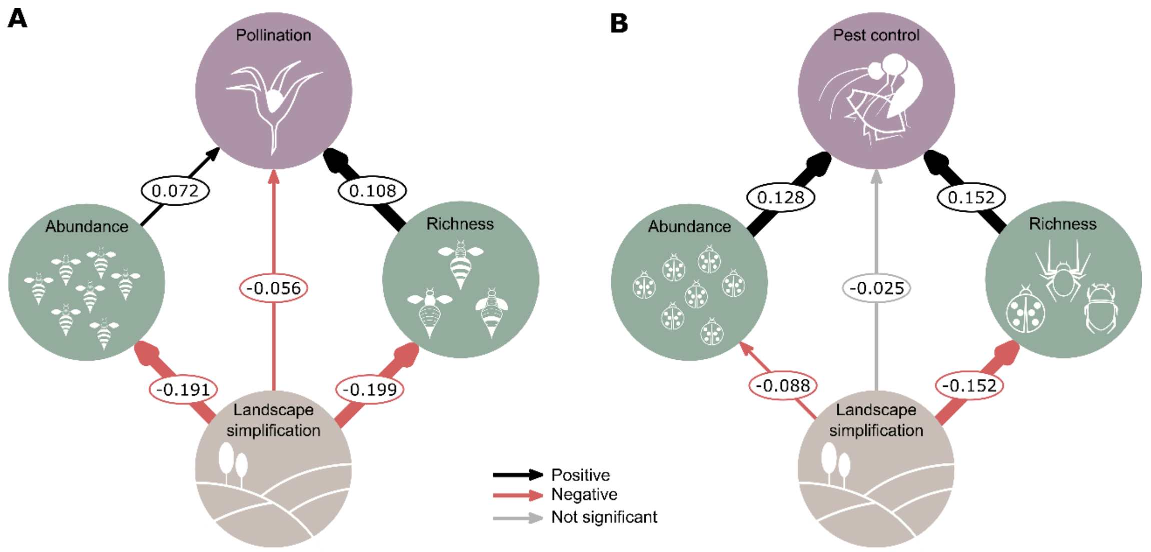


**Fig. S4.**

**Direct and indirect landscape simplification effects on ecosystem services via changes in richness and abundance.** (**A**) Path model representing direct and indirect effects of landscape simplification on pollination through changes in pollinator richness and abundance (*N* = 821 fields of 52 crop systems. (**B**) Path model representing direct and indirect effects of landscape simplification on pest control services through changes in natural enemy richness and abundance (*N* = 654 fields of 37 crop systems). Path coefficients are effect sizes estimated from the median of the posterior distribution of the model. Black and red arrows represent positive or negative effects, respectively. Arrow widths are proportional to highest density intervals (HDIs). Grey arrows represent non-evident effects (HDIs overlapped zero).

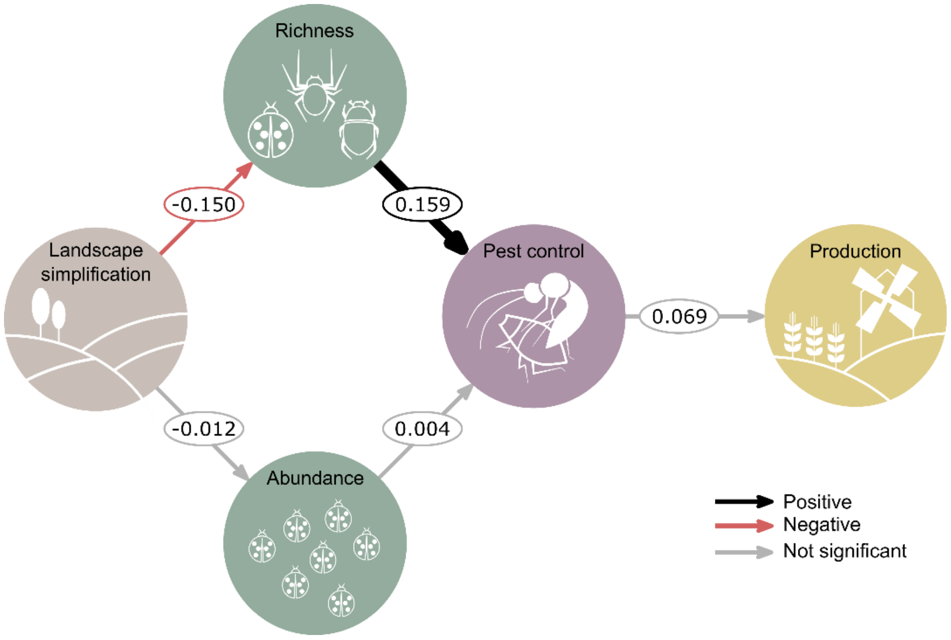


**Fig. S5.**

**Direct and cascading landscape simplification effects on final crop production via changes in natural enemy richness, abundance and pest control (all sites together, with and without insecticide application).** Path coefficients are effect sizes estimated from the median of the posterior distribution of the model ($N$ = 236 fields of 15 crop systems). Black and red arrows represent positive or negative effects, respectively. Arrow widths are proportional to highest density intervals (HDIs). Grey arrows represent non-significant effects (HDIs overlapped zero).

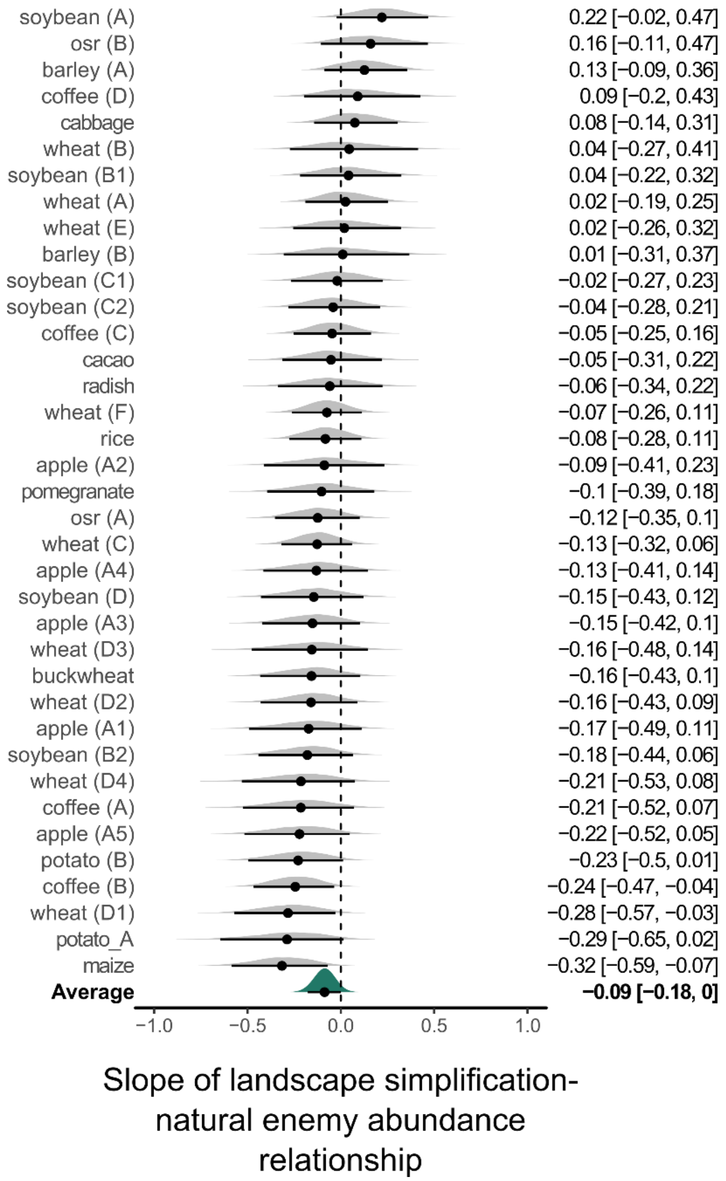


**Fig. S6.**

**Forest plot of the effect of landscape simplification on natural enemy abundance for individual crop systems.** Each posterior distribution represents medians (symbol centres) and 90% density intervals (black lines).

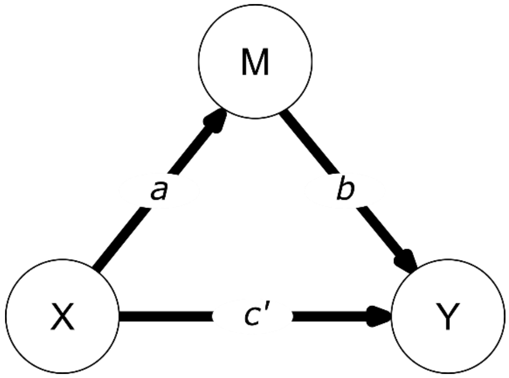


**Fig. S7.**

**Mediation model.** Mediation analysis is a statistical procedure to test whether the effect of an independent variable X on a dependent variable Y (X → Y) is at least partly explained via the inclusion of a third hypothetical variable, the mediator variable M (X → M → Y). The three causal paths *a*, *b*, and *c'* represent X's effect on M, M's effect on Y, and X's effect on Y while accounting for M, respectively. The three causal paths correspond to parameters from two regression models, one in which M is the outcome and X the predictor, and one in which Y is the outcome and X and M the simultaneous predictors.

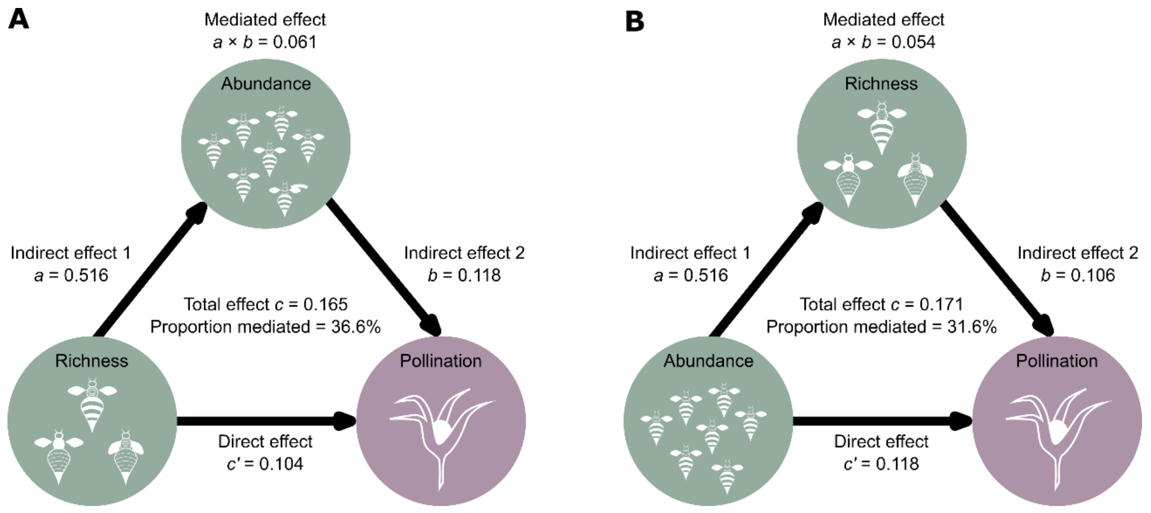


**Fig. S8.**

**Direct and mediated effects of pollinator richness and abundance (with honey bees) on pollination.** (**A**) Path model of pollinator richness as a predictor of pollination, mediated by pollinator abundance. (**B**) Path model of pollinator abundance as a predictor of pollination, mediated by pollinator richness. $N$ = 821 fields of 52 crop systems. Coefficients of the three causal paths ($a$, $b$, $c'$) correspond to the median of the posterior distribution of the model. The proportion mediated is the mediated effect ($a \times b$) divided by the total effect ($c$).

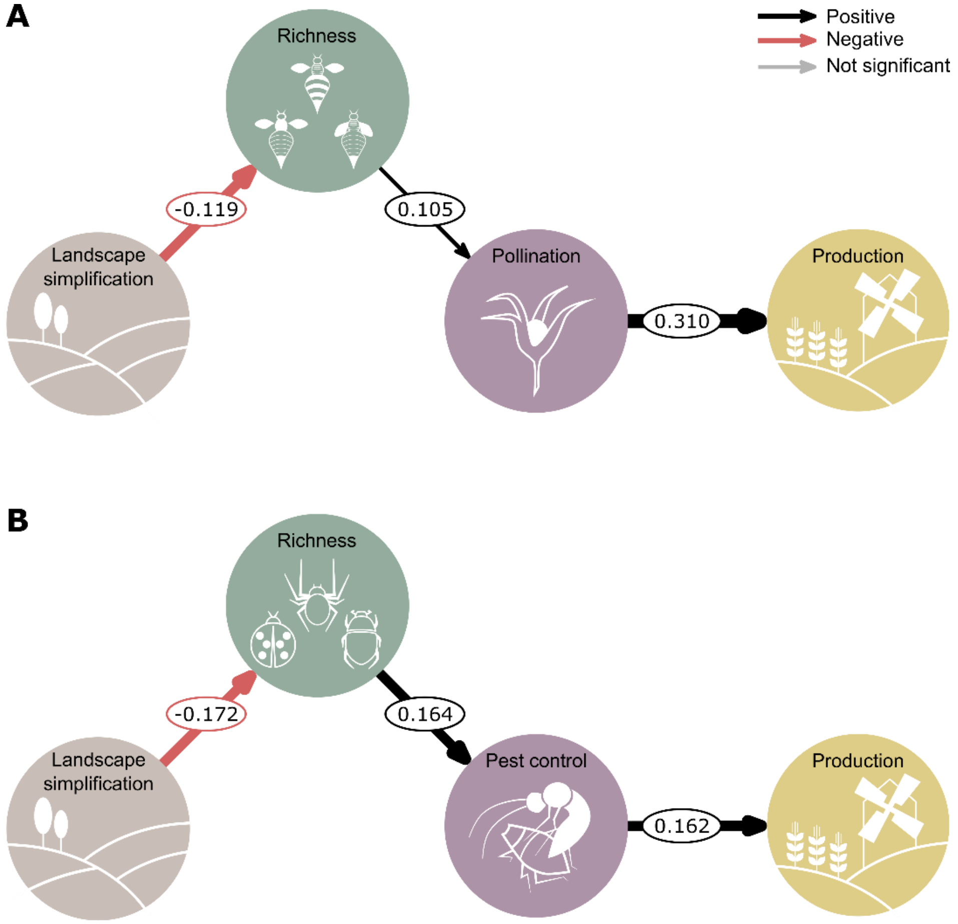


**Fig. S9.**

**Direct and cascading landscape simplification effects on area-based yield via changes in richness, abundance and ecosystem services**. (**A**) Path model representing direct and indirect effects of landscape simplification on final area-based yield through changes in pollinator richness, abundance and pollination ($N$ = 203 fields of 13 crop systems). (**B**) Path model representing direct and indirect effects of landscape simplification on final area-based yield through changes in natural enemy richness, abundance and pest control ($N$ = 102 fields of 7 crop systems). Path coefficients are effect sizes estimated from the median of the posterior distribution of the model. Black and red arrows represent positive or negative effects, respectively. Arrow widths are proportional to highest density intervals (HDIs). Grey arrows represent non-significant effects (HDIs overlapped zero).

**Table S1.**
**List of 89 crop systems considered in our analyses**.

| Crop and system code | Reference and (or) data holder contact | Crop species | Country, region | Study year | Sites (with yield) | Sampling methods | Taxa | Functions | Production |
|---|---|---|---|---|---|---|---|---|---|
| **Pollination studies** | | | | | | | | | |
| acerola | (*30*) Freitas, | *Malpighia emarginata* | Brazil, Ceará | 2011 | 8 | active | bees | fruit set | - |
| apple (A) | (*58*) Boreux, | *Malus domestica* | Germany, Lake Constance | 2015 | 25 | active | bees | fruit set | - |
| apple (B) | (*59*) Garratt, | *Malus domestica* | UK, Kent | 2011 | 8 | active, passive | bees | fruit set | - |
| apple (C1) | (*60*) de Groot, | *Malus domestica* | Netherlands, Betuwe | 2013 | 8 (4) | active | bees, hoverflies | fruit set | crop yield |
| apple (C2) | (*60*) de Groot, | *Malus domestica* | Netherlands, Betuwe | 2014 | 10 (9) | active | bees, hoverflies | fruit set | crop yield |
| apple (D1) | (*61*) Mallinger, | *Malus domestica* | USA, Wisconsin | 2012 | 17 | passive | bees | fruit set | - |
| apple (D2) | (*61*) Mallinger, | *Malus domestica* | USA, Wisconsin | 2013 | 19 | passive | bees | fruit set | - |
| bean (A) | Ekroos, | *Vicia faba* | Sweden, Scania | 2016 | 16 (16) | active | bees | seed set | plant yield |
| bean (B) | (*62*) Garratt, | *Vicia faba* | UK, Berkshire | 2011 | 8 | active, passive | bees | seed set | - |
| bean (C) | Ramos, Silva | *Phaseolus vulgaris* | Brazil, Goias/DF | 2015/2016 | 22 (22) | active | bees | seed set | crop yield |
| blueberry (A) | Cavigliasso, | *Vaccinium corymbosum* | Argentina, Espinal-Ñandubay | 2016 | 13 | active | bees, wasps, hoverflies | fruit set | - |
| blueberry (B1) | (*60*) de Groot, | *Vaccinium corymbosum* | Netherlands, Limburg/Overijssel | 2013 | 10 (9) | active | bees | fruit set | crop yield |
| blueberry (B2) | (*60*) de Groot, | *Vaccinium corymbosum* | Netherlands, Limburg/Overijssel | 2014 | 15 (13) | active | bees | fruit set | crop yield |
| buckwheat (A1) | (*63*, *64*) Taki, | *Fagopyrum esculentum* | Japan, Ibaraki | 2007 | 15 | active | bees, butterflies, flies, wasps | seed set | - |
| buckwheat (A2) | (*63*, *64*) Taki, | *Fagopyrum esculentum* | Japan, Ibaraki | 2008 | 17 | active | bees, butterflies, flies, wasps | seed set | - |
| cashew | (*14*) Freitas, | *Anacardium occidentale* | Brazil, Ceará | 2012 | 10 (10) | active | bees | fruit set | crop yield |

| | | | | | | | | | |
|---|---|---|---|---|---|---|---|---|---|
| cherry | (*65*) Holzschuh, | *Prunus avium* | Germany, Hesse | 2008 | 7 | active | bees | fruit set | - |
| coffee (A) | (*33*, *66*, *67*) Boreux, | *Coffea canephora* | India, Kodagu | 2008 | 53 (51) | active | bees | fruit set | plant yield |
| coffee (B) | (*68*) Classen, | *Coffea arabica* | Tanzania, Kilimanjaro | 2011/2012 | 11 (7) | active, passive | bees | fruit set | plant yield |
| coffee (C) | (*69*) Hipólito, | *Coffea arabica* | Brazil, Chapada Diamantina | 2013 | 30 (28) | active | bee, flies, butterflies, beetles, wasps | fruit set | crop yield |
| coffee (D1) | (*66*, *70*, *71*) Krishnan, | *Coffea canephora* | India, Kodagu | 2007 | 35 | active | bees | fruit set | - |
| coffee (D2) | (*66*, *70*, *71*) Krishnan, | *Coffea canephora* | India, Kodagu | 2008 | 37 | active | bees | fruit set | |
| coffee (E) | (*72*) Krishnan, Nesper, | *Coffea canephora* | India, Kodagu | 2014 | 49 (49) | active | bees | fruit set | crop yield |
| cotton | (*73*) Cusser, | *Gossypium hirsutum* | USA, Gulf Coast Texas | 2014 | 11 | active | bee, hoverflies, butterflies, beetles | fruit set | - |
| grapefruit | (*74*, *75*) Chacoff, | *Citrus paradisi* | Argentina, Yungas | 2000 | 6 | active | bee, flies, butterflies, wasps | fruit set | |
| leek (A) | (*34*) Fijen, | *Allium porrum* | France, Loire | 2016 | 18 (18) | active | bees, wasps, hoverflies | seed set | plant yield |
| leek (B) | (*34*) Fijen, | *Allium porrum* | Italy, South Italy | 2016 | 18 (18) | active | bees, wasps, hoverflies | seed set | plant yield |
| lemon | Chacoff, | *Citrus limon* | Argentina, Yungas | 2015 | 9 | active | bee, flies, butterflies, wasps | fruit set | - |
| mango (A) | (*76*) Carvalheiro, | *Mangifera indica* | South Africa, Limpopo | 2008 | 8 | active | bee, flies, butterflies, beetles, wasps | fruit set | - |
| mango (B) | (*77*) Carvalheiro, | *Mangifera indica* | South Africa, Limpopo | 2009 | 14 (10) | active | bee, flies, butterflies, beetles, wasps | fruit set | plant yield |
| mango (C) | Rader, | *Mangifera indica* | Australia, Queensland | 2014 | 10 | active | bees, flies, hoverflies, beetles, moths, butterflies | fruit set | - |
| mango (D) | Willcox, | *Mangifera indica* | Australia, Queensland | 2016 | 7 | active | bees, flies, hoverflies, beetles, | fruit set | - |

| | | | | | | | | | |
|---|---|---|---|---|---|---|---|---|---|
| | | | | | | | moths, butterflies | | |
| osr (A) | Andersson, | *Brassica napus* | Sweden, Scania | 2010 | 6 | active | bees, hoverflies | seed set | - |
| osr (B) | (*35*) Bartomeus, Gagic, | *Brassica napus* | Sweden, Västergötland | 2013 | 12 (9) | active | bees, butterflies | seed set | crop yield |
| osr (C) | (*62*) Garratt, | *Brassica napus* | UK, Yorkshire | 2012 | 8 | active, passive | bees | seed set | - |
| osr (D) | (*78, 79*) Stanley, | *Brassica napus* | Ireland, South-East | 2010 | 3 | active | bees, hoverflies | seed set | - |
| osr (E) | Sutter, | *Brassica napus* | Switzerland, Zurich | 2014 | 18 (18) | active | bees, hoverflies | seed set | crop yield |
| osr (F) | (*80*) Zou Yi, | *Brassica napus* | China, Jiangxi | 2015 | 18 | passive | bees, hoverflies, butterflies | fruit set | - |
| raspberry | (*29*) Saez, | *Rubus idaeus* | Argentina, Comarca Andina | 2014 | 16 (16) | active | bees | fruit set | crop yield |
| red clover | Rundlöf, | *Trifolium pratense* | Sweden, Scania | 2013 | 6 (6) | active | bees | seed set | crop yield |
| strawberry (A) | Andersson, | *Fragaria* × *ananassa* | Sweden, Scania | 2009 | 11 | passive | bees, hoverflies | fruit set | - |
| strawberry (B) | Baensch, Tscharntke, Westphal, | *Fragaria* × *ananassa* | Germany, Lower Saxony, | 2015 | 8 (8) | active | bees | Δ fruit weight | plant yield |
| strawberry (C1) | (*81*) Grab, | *Fragaria* × *ananassa* | USA, New York | 2012 | 11 (11) | active, passive | bees | Δ fruit weight | plant yield |
| strawberry (C2) | Grab, | *Fragaria* × *ananassa* | USA, New York | 2014 | 27 (27) | active | bees | seed set | plant yield |
| strawberry (C3) | (*82*) Grab, | *Fragaria* × *ananassa* | USA, New York | 2015 | 14 (14) | active | bees | seed set | plant yield |
| strawberry (D) | Garratt, | *Fragaria* × *ananassa* | UK, Yorkshire | 2011 | 7 (7) | active, passive | bees | Δ fruit weight | plant yield |
| strawberry (E) | Klatt, | *Fragaria* × *ananassa* | Germany, Lower Saxony | 2010 | 8 (8) | active | bees | fruit set | plant yield |
| strawberry (F) | Krewenka, | *Fragaria* × *ananassa* | Germany, Lower Saxony | 2005 | 10 (10) | active | bees | fruit set | crop yield |
| strawberry (G) | Sciligo, | *Fragaria* × *ananassa* | USA, California | 2012 | 15 (15) | active, passive | bees | Δ fruit weight | plant yield |
| strawberry (H) | (*83*) Stewart, | *Fragaria* × *ananassa* | Sweden, Scania | 2014 | 27 (27) | active | hoverflies | fruit set | plant yield |
| sunflower (A) | (*32*) Carvalheiro, | *Helianthus annuus* | South Africa, Limpopo | 2009 | 28 | active | bee, flies, butterflies , beetles, wasps | seed set | - |
| sunflower (B) | Scheper, | *Helianthus annuus* | France, Poitou-Charentes | 2015 | 24 | active | bees, hoverflies | seed set | - |

| **Pest control studies** | | | | | | | | | |
|---|---|---|---|---|---|---|---|---|---|
| apple (A1) | (*84*, *85*) Lavigne, | *Malus domestica* | France, Provence-Alpes-Côte d'Azur | 2006 | 9 | active | parasitoids | sentinel exp. (enemy activity) | - |
| apple (A2) | (*84*, *85*) Lavigne, | *Malus domestica* | France, Provence-Alpes-Côte d'Azur | 2007 | 6 | active | parasitoids | sentinel exp. (enemy activity) | - |
| apple (A3) | (*84*, *85*) Lavigne, | *Malus domestica* | France, Provence-Alpes-Côte d'Azur | 2008 | 17 | active | parasitoids | sentinel exp. (enemy activity) | - |
| apple (A4) | (*84*, *85*) Lavigne, | *Malus domestica* | France, Provence-Alpes-Côte d'Azur | 2009 | 12 | active | parasitoids | sentinel exp. (enemy activity) | - |
| apple (A5) | (*84*, *85*) Lavigne, | *Malus domestica* | France, Provence-Alpes-Côte d'Azur | 2010 | 14 | active | parasitoids | sentinel exp. (enemy activity) | - |
| barley (A) | (*86*) Caballero-Lopez, | *Hordeum vulgare* | Sweden, Scania | 2007 | 20 | active, passive | carabids, ladybugs, parasitoids | sentinel exp. (enemy activity) | - |
| barley (B) | (*87*–*89*) Tamburini, | *Hordeum vulgare* | Italy, Friuli Venezia-Giulia | 2014 | 5 (5) | passive | carabids | cage exp. (infestation) | crop yield |
| buckwheat | (*90*) Taki, | *Fagopyrum esculentum* | Japan, Ibaraki | 2008 | 15 | passive | ladybugs, lacewings | sentinel exp. (enemy activity) | - |
| cabbage | (*91*) Letourneau, | *Brassica oleracea* | USA, Monterey Bay Area | 2006 | 33 | passive | parasitoids | sentinel exp. (enemy activity) | - |
| cacao | (*92*) Maas, | *Theobroma cacao* | Indonesia, Sulawesi | 2010 | 15 (15) | active | spiders | cage exp. (crop damage) | plant yield |
| coffee (A) | Schleuning, Schmack, | *Coffea arabica* | Tanzania, Kilimanjaro | 2011/2012 | 11 (6) | passive | bats, birds | cage exp. (crop damage) | plant yield |
| coffee (B) | Iverson, | *Coffea arabica* | Mexico, Soconusco | 2012 | 37 (35) | passive | parasitoids | sentinel exp. (enemy activity) | crop yield |
| coffee (C) | Iverson, | *Coffea arabica* | Puerto Rico, Utuado | 2013 | 36 | passive | parasitoids, wasps | cage exp. (crop damage) | - |
| coffee (D) | Martinez-Salinas, | *Coffea arabica* | Costa Rica, Turrialba | 2013 | 10 | passive | birds | cage exp. (crop damage) | - |
| maize | (*93*) O'Rourke, | *Zea mays* | USA, New York | 2006 | 26 | passive | ladybugs | pest damage | - |
| osr (A) | (*94*) Jonsson, | *Brassica napus* | New Zealand, Canterbury | 2007 | 26 | active | hoverflies, ladybugs, lacewings | pest damage | - |
| osr (B) | Sutter, | *Brassica napus* | Switzerland, Zurich | 2014 | 18 (18) | passive | carabids | sentinel exp. (enemy activity) | crop yield |

| | | | | | | | | | |
|---|---|---|---|---|---|---|---|---|---|
| pomegranate | (*95*) Keasar, | *Punica granatum* | Israel, Hefer Valley | 2014 | 10 | active | spiders, parasitoids | pest damage | - |
| potato (A) | (*96*) Martin, | *Solanum tuberosum* | South Korea, Haean | 2009 | 6 (2) | active, passive | birds, carabids, hoverflies, parasitoids, rove beetles, wasps | pest damage | plant yield |
| potato (B) | (*97*) Poveda, | *Solanum tuberosum* | Colombia, Cundinamarca | 2007 | 11 (11) | active, passive | carabids, hoverflies, ladybugs, lacewings, parasitoids | pest damage | crop yield |
| radish | (*96*) Martin, | *Raphanus raphanistrum* subsp. *sativus* | South Korea, Haean | 2009 | 8 (5) | active, passive | birds, carabids, hoverflies, parasitoids, rove beetles, wasps | pest damage | plant yield |
| rice | (*98*) Takada, | *Oryza sativa* | Japan, Miyagi | 2008 | 44 | active | spiders | pest damage | - |
| soybean (A) | (*25*) Kim, | *Glycine max* | USA, Upper Midwest | 2012 | 35 (33) | passive | flower bugs, ladybugs | cage exp. (infestation) | plant yield |
| soybean (B1) | (*99*) Mitchell, | *Glycine max* | Canada, Montérégie | 2010 | 15 (15) | active | hoverflies, ladybugs, lacewings, true bugs | pest damage | crop yield |
| soybean (B2) | (*99*) Mitchell, | *Glycine max* | Canada, Montérégie | 2011 | 19 (19) | active | hoverflies, ladybugs, lacewings, true bugs | pest damage | crop yield |
| soybean (C1) | Molina, | *Glycine max* | Argentina, North Buenos Aires | 2011 | 20 | active | parasitoids | sentinel exp. (enemy activity) | - |
| soybean (C2) | Molina, | *Glycine max* | Argentina, North Buenos Aires | 2012 | 20 | active | parasitoids | sentinel exp. (enemy activity) | - |
| soybean (D) | (*96*) Martin, | *Glycine max* | South Korea, Haean | 2009 | 8 (6) | active, passive | birds, carabids, hoverflies, parasitoids, rove beetles, wasps | pest damage | plant yield |

| | | | | | | | | | |
|---|---|---|---|---|---|---|---|---|---|
| wheat (A) | (*100*) Bommarco, | *Triticum aestivum* | Sweden, Scania | 2007 | 31 (31) | passive | carabids | sentinel exp. (enemy activity) | crop yield |
| wheat (B) | (*86*) Caballero-Lopez, | *Triticum aestivum* | Sweden, Scania | 2007 | 4 | active, passive | carabids, ladybugs, parasitoid s | sentinel exp. (enemy activity) | - |
| wheat (C) | Kim, | *Triticum aestivum* | USA, Upper Midwest | 2012 | 24 (24) | active, passive | flower bugs, ladybugs | cage exp. (infestation) | plant yield |
| wheat (D1) | (*101*) Plećaš, | *Triticum aestivum* | Serbia, Pacevacki Rit | 2008 | 18 | active | parasitoid s | sentinel exp. (enemy activity) | - |
| wheat (D2) | (*101*) Plećaš, | *Triticum aestivum* | Serbia, Pacevacki Rit | 2009 | 17 | active | parasitoid s | sentinel exp. (enemy activity) | - |
| wheat (D3) | (*101*) Plećaš, | *Triticum aestivum* | Serbia, Pacevacki Rit | 2010 | 8 | active | parasitoid s | sentinel exp. (enemy activity) | - |
| wheat (D4) | (*101*) Plećaš, | *Triticum aestivum* | Serbia, Pacevacki Rit | 2011 | 10 | active | parasitoid s | sentinel exp. (enemy activity) | - |
| wheat (E) | (*87–89*) Tamburini, | *Triticum aestivum* | Italy, Friuli Venezia-Giulia | 2014 | 11 (11) | passive | carabids | cage exp. (infestation) | crop yield |
| wheat (F) | (*102*) Tschumi, | *Triticum aestivum* | Switzerland, Central Plateau | 2012 | 25 | active, passive | carabids, ladybugs, true bugs | pest damage | - |

**Table S2.**

**Model output for richness-ecosystem service relationships.** (**A**) Richness was calculated as the number of unique taxa sampled per crop system. (**B**) Richness was calculated considering only organisms classified at the fine taxonomy level (i.e. species- or morphospecies-levels). Posterior samples were summarized based on the Bayesian point estimate (median), standard error (median absolute deviation), and 80%, 90% and 95% highest density intervals (HDIs). HDIs that do not include zero are reported in bold.

(**A**)

| Parameter | Estimate | SE | HDI (80%) | HDI (90%) | HDI (95%) |
|---|---|---|---|---|---|
| ***Pollination*** | | | | | |
| Intercept | 0.0004 | 0.0311 | [-0.0419, 0.0411] | [-0.0531, 0.0544] | [-0.0670, 0.0622] |
| Pollinator richness | 0.1532 | 0.0353 | **[0.1062, 0.1962]** | **[0.0951, 0.2110]** | **[0.0865, 0.2266]** |
| ***Pest control*** | | | | | |
| Intercept | -0.0003 | 0.0353 | [-0.0434, 0.0485] | [-0.0579, 0.0589] | [-0.0724, 0.0657] |
| Natural enemy richness | 0.2132 | 0.0412 | **[0.1585, 0.2646]** | **[0.1451, 0.2810]** | **[0.1314, 0.2954]** |

(**B**)

| Parameter | Estimate | SE | HDI (80%) | HDI (90%) | HDI (95%) |
|---|---|---|---|---|---|
| ***Pollination*** | | | | | |
| Intercept | 0.0010 | 0.0333 | [-0.0409, 0.0421] | [-0.0536, 0.0537] | [-0.0662, 0.0617] |
| Pollinator richness | 0.1535 | 0.0356 | **[0.1096, 0.2006]** | **[0.0967, 0.2141]** | **[0.0848, 0.2256]** |
| ***Pest control*** | | | | | |
| Intercept | 0.0001 | 0.0401 | [-0.0536, 0.0514] | [-0.0712, 0.0646] | [-0.0834, 0.0775] |
| Natural enemy richness | 0.2262 | 0.0484 | **[0.1671, 0.2913]** | **[0.1420, 0.3022]** | **[0.1315, 0.3225]** |

**Table S3.**

**Model output for path models testing direct and indirect effects (mediated by changes in abundance) of richness on ecosystem services.** Posterior samples were summarized based on the Bayesian point estimate (median), standard error (median absolute deviation), and 80%, 90% and 95% highest density intervals (HDIs). HDIs that do not include zero are reported in bold.

| Effect | Estimate | SE | HDI (80%) | HDI (90%) | HDI (95%) |
|---|---|---|---|---|---|
| ***Pollination*** | | | | | |
| Richness → Pollination | 0.1058 | 0.0428 | **[0.0511, 0.1635]** | **[0.0326, 0.1779]** | **[0.0199, 0.1933]** |
| Richness → Abundance | 0.5701 | 0.0379 | **[0.5222, 0.6212]** | **[0.5044, 0.6319]** | **[0.4900, 0.6449]** |
| Abundance → Pollination | 0.0804 | 0.0460 | **[0.0232, 0.1401]** | **[0.0057, 0.1564]** | [-0.0140, 0.1665] |
| *Causal mediation analysis* | | | | | |
| Direct effect | 0.1058 | | **[0.0511, 0.1635]** | **[0.0326, 0.1779]** | **[0.0199, 0.1933]** |
| Indirect effect | 0.0456 | | **[0.0142, 0.0812]** | **[0.0038, 0.0903]** | [-0.0079, 0.0955] |
| Total effect | 0.1512 | | **[0.1036, 0.1997]** | **[0.0905, 0.2136]** | **[0.0774, 0.2239]** |
| Proportion mediated | 30.1% | | | | |
| ***Pest control*** | | | | | |
| Richness → Pest control | 0.1413 | 0.0434 | **[0.0832, 0.1951]** | **[0.0684, 0.2105]** | **[0.0564, 0.2275]** |
| Richness → Abundance | 0.4447 | 0.0494 | **[0.3782, 0.5070]** | **[0.3646, 0.5315]** | **[0.3467, 0.5452]** |
| Abundance → Pest control | 0.1481 | 0.0553 | **[0.0772, 0.2170]** | **[0.0612, 0.2406]** | **[0.0467, 0.2619]** |
| *Causal mediation analysis* | | | | | |
| Direct effect | 0.1413 | | **[0.0832, 0.1951]** | **[0.0684, 0.2105]** | **[0.0564, 0.2275]** |
| Indirect effect | 0.0650 | | **[0.0306, 0.0986]** | **[0.0242, 0.1119]** | **[0.0175, 0.1226]** |
| Total effect | 0.2084 | | **[0.1545, 0.2629]** | **[0.1398, 0.2778]** | **[0.1276, 0.2945]** |
| Proportion mediated | 31.2% | | | | |

**Table S4.**

**Model output for path models testing direct and indirect effects (mediated by changes in richness) of abundance on ecosystem services.** Posterior samples were summarized based on the Bayesian point estimate (median), standard error (median absolute deviation), and 80%, 90% and 95% highest density intervals (HDIs). HDIs that do not include zero are reported in bold.

| Effect | Estimate | SE | HDI (80%) | HDI (90%) | HDI (95%) |
|---|---|---|---|---|---|
| ***Pollination*** | | | | | |
| Abundance → Pollination | 0.0807 | 0.0455 | **[0.0276, 0.1438]** | **[0.0045, 0.1534]** | [-0.0077, 0.1728] |
| Abundance → Richness | 0.5706 | 0.0376 | **[0.5215, 0.6206]** | **[0.5053, 0.6342]** | **[0.4915, 0.6466]** |
| Richness → Pollination | 0.1052 | 0.0432 | **[0.0479, 0.1599]** | **[0.0362, 0.1799]** | **[0.0171, 0.1901]** |
| *Causal mediation analysis* | | | | | |
| Direct effect | 0.0807 | | **[0.0276, 0.1438]** | **[0.0045, 0.1534]** | [-0.0077, 0.1728] |
| Indirect effect | 0.0597 | | **[0.0267, 0.0912]** | **[0.0184, 0.1020]** | **[ 0.0117, 0.1119]** |
| Total effect | 0.1409 | | **[0.0900, 0.1874]** | **[0.0753, 0.2012]** | **[ 0.0665, 0.2172]** |
| Proportion mediated | 42.4% | | | | |
| ***Pest control*** | | | | | |
| Abundance → Pest control | 0.1409 | 0.0522 | **[0.0677, 0.2060]** | **[0.0548, 0.2353]** | **[0.0366, 0.2538]** |
| Abundance → Richness | 0.4396 | 0.0522 | **[0.3762, 0.5058]** | **[0.3566, 0.5239]** | **[0.3404, 0.5414]** |
| Richness → Pest control | 0.1495 | 0.0430 | **[0.0946, 0.2058]** | **[0.0770, 0.2195]** | **[0.0655, 0.2351]** |
| *Causal mediation analysis* | | | | | |
| Direct effect | 0.1409 | | **[0.0677, 0.2060]** | **[0.0548, 0.2353]** | **[0.0366, 0.2538]** |
| Indirect effect | 0.0651 | | **[0.0398, 0.0922]** | **[0.0324, 0.0997]** | **[0.0280, 0.1088]** |
| Total effect | 0.2069 | | **[0.1405, 0.2688]** | **[0.1227, 0.2891]** | **[0.1133, 0.3135]** |
| Proportion mediated | 31.5% | | | | |

**Table S5.**

**Model output for path models testing direct and indirect effects (mediated by changes in richness) of landscape simplification on ecosystem services.** Posterior samples were summarized based on the Bayesian point estimate (median), standard error (median absolute deviation), and 80%, 90% and 95% highest density intervals (HDIs). HDIs that do not include zero are reported in bold.

| Effect | Estimate | SE | HDI (80%) | HDI (90%) | HDI (95%) |
|---|---|---|---|---|---|
| ***Pollination*** | | | | | |
| Landscape → Pollination | -0.0573 | 0.0409 | **[-0.1083, -0.0041]** | [-0.1203, 0.0147] | [-0.1374, 0.0229] |
| Landscape → Richness | -0.1984 | 0.0453 | **[-0.2593, -0.1430]** | **[-0.2750, -0.1263]** | **[-0.2909, -0.1119]** |
| Richness → Pollination | 0.1543 | 0.0362 | **[0.1060, 0.1992]** | **[0.0937, 0.2148]** | **[0.0815, 0.2278]** |
| *Causal mediation analysis* | | | | | |
| Direct effect | -0. 0573 | | **[-0.1083, -0.0041]** | [-0.1203, 0.0147] | [-0.1374, 0.0229] |
| Indirect effect | -0.0293 | | **[-0.0425, -0.0168]** | **[-0.0465, -0.0136]** | **[-0.0515, -0.0117]** |
| Total effect | -0.0859 | | **[-0.1391, -0.0361]** | **[-0.1560, -0.0239]** | **[-0.1642, -0.0074]** |
| Proportion mediated | 34.0% | | | | |
| | | | | | - |
| ***Pest control*** | | | | | |
| Landscape → Pest control | -0.0285 | 0.0442 | [-0.0864, -0.0289] | [-0.1043, 0.0461] | [-0.1248, 0.0570] |
| Landscape → Richness | -0.1510 | 0.0479 | **[-0.2123, -0.0886]** | **[-0.2299, -0.0706]** | **[-0.2491, -0.0581]** |
| Richness → Pest control | 0.2114 | 0.0418 | **[0.1609, 0.2682]** | **[0.1429, 0.2810]** | **[0.1315, 0.2962]** |
| *Causal mediation analysis* | | | | | |
| Direct effect | -0.0285 | | [-0.0864, -0.0289] | [-0.1043, 0.0461] | [-0.1248, 0.0570] |
| Indirect effect | -0.0311 | | **[-0.0460 -0.0149]** | **[-0.0523, -0.0118]** | **[-0.0578, -0.0083]** |
| Total effect | -0.0610 | | **[-0.1214, -0.0060]** | [-0.1378, 0.0120] | [-0.1511, 0.0301] |
| Proportion mediated | 50.9% | | | | |

**Table S6.**

**Model output for path models testing the direct and cascading landscape simplification effects on ecosystem services via changes in richness and abundance.** Posterior samples were summarized based on the Bayesian point estimate (median), standard error (median absolute deviation), and 80%, 90% and 95% highest density intervals (HDIs). HDIs that do not include zero are reported in bold.

| Effect | Estimate | SE | HDI (80%) | HDI (90%) | HDI (95%) |
|---|---|---|---|---|---|
| ***Pollination*** | | | | | |
| Landscape → Richness | -0.1991 | 0.0458 | **[-0.2593, -0.1431]** | **[-0.2779, -0.1269]** | **[-0.2918, -0.1109]** |
| Landscape → Abundance | -0.1914 | 0.0462 | **[-0.2503, -0.1302]** | **[-0.2721, -0.1167]** | **[-0.2812, -0.0955]** |
| Landscape → Pollination | -0.0559 | 0.0390 | **[-0.1043, -0.0035]** | [-0.1223, 0.0078] | [-0.1351, 0.0215] |
| Richness → Pollination | 0.1082 | 0.0430 | **[ 0.0517, 0.1629]** | **[ 0.0366 0.1810]** | **[ 0.0197, 0.1924]** |
| Abundance → Pollination | 0.0721 | 0.0444 | **[0.0155, 0.1313]** | [-0.0017, 0.1479] | [-0.0229, 0.1558] |
| ***Pest control*** | | | | | |
| Landscape → Richness | -0.1515 | 0.0471 | **[-0.2160, -0.0939]** | **[-0.2322, -0.0730]** | **[-0.2430, -0.0471]** |
| Landscape → Abundance | -0.0880 | 0.0511 | **[-0.1617, -0.0240]** | [-0.1727, 0.0044] | [-0.1968, 0.0148] |
| Landscape → Pest control | -0.0250 | 0.0436 | [-0.0785, 0.0316] | [-0.0971, 0.0451] | [-0.1128, 0.0559] |
| Richness → Pest control | 0.1524 | 0.0436 | **[0.0928, 0.2049]** | **[0.0822, 0.2272]** | **[0.0642, 0.2385]** |
| Abundance → Pest control | 0.1282 | 0.0540 | **[0.0597, 0.1967]** | **[0.0398, 0.2146]** | **[0.0323, 0.2403]** |

**Table S7.**

**Model output for path models testing the direct and cascading landscape simplification effects on final crop production via changes in richness, abundance and ecosystem services.** Posterior samples were summarized based on the Bayesian point estimate (median), standard error (median absolute deviation), and 80%, 90% and 95% highest density intervals (HDIs). HDIs that do not include zero are reported in bold.

| Effect | Estimate | SE | HDI (80%) | HDI (90%) | HDI (95%) |
|---|---|---|---|---|---|
| ***Pollination*** | | | | | |
| Landscape → Richness | -0.1870 | 0.0571 | **[-0.2631, -0.1128]** | **[-0.2876, -0.0926]** | **[-0.3067, -0.0722]** |
| Landscape → Abundance | -0.1988 | 0.0542 | **[-0.2680, -0.1270]** | **[-0.2915, -0.1069]** | **[-0.3103, -0.0884]** |
| Richness → Pollination | 0.1477 | 0.0638 | **[0.0645, 0.2281]** | **[0.0485, 0.2568]** | **[0.0212, 0.2712]** |
| Abundance → Pollination | 0.0104 | 0.0667 | [-0.0742, 0.0961] | [-0.0951, 0.1243] | [-0.1250, 0.1373] |
| Pollination → Production | 0.3388 | 0.0868 | **[0.2268, 0.4509]** | **[0.1910, 0.4813]** | **[0.1549, 0.5070]** |
| ***Pest control*** | | | | | |
| Landscape → Richness | -0.2073 | 0.0840 | **[-0.3197, -0.0977]** | **[-0.3502, -0.0554]** | **[-0.3915, -0.0216]** |
| Landscape → Abundance | -0.0304 | 0.1060 | [-0.1759, 0.1106] | [-0.2242, 0.1587] | [-0.2745, 0.1938] |
| Richness → Pest control | 0.2255 | 0.0786 | **[0.1201, 0.3236]** | **[0.0932, 0.3573]** | **[0.0730, 0.3905]** |
| Abundance → Pest control | 0.0040 | 0.0793 | [-0.1016, 0.1064] | [-0.1331, 0.1413] | [-0.1572, 0.1769] |
| Pest control → Production | 0.1395 | 0.0786 | **[0.0404, 0.2451]** | **[0.0151 0.2822]** | [-0.0257, 0.3011] |

**Table S8.**

**Model output for path models testing direct and mediated effects of pollinator richness and abundance (with honey bees) on pollination.** Posterior samples were summarized based on the Bayesian point estimate (median), standard error (median absolute deviation), and 80%, 90% and 95% highest density intervals (HDIs). HDIs that do not include zero are reported in bold.

| Effect | Estimate | SE | HDI (80%) | HDI (90%) | HDI (95%) |
|---|---|---|---|---|---|
| ***Pollination*** | | | | | |
| Richness → Pollination | 0. 1043 | 0.0419 | **[0.0524, 0.1588]** | **[0.0356, 0.1715]** | **[0.0250, 0.1878]** |
| Richness → Abundance | 0.5160 | 0.0377 | **[0.4682, 0.5659]** | **[0.4511, 0.5784]** | **[0.4328, 0.5882]** |
| Abundance → Pollination | 0.1183 | 0.0452 | **[0.0617, 0.1768]** | **[0.0430, 0.1903]** | **[0.0278, 0.2038]** |
| *Causal mediation analysis* | | | | | |
| Direct effect | 0.1043 | | **[0.0524, 0.1588]** | **[0.0356, 0.1715]** | **[0.0250, 0.1878]** |
| Indirect effect | 0.0606 | | **[0.0292, 0.0893]** | **[0.0226, 0.0996]** | **[0.0162, 0.1085]** |
| Total effect | 0.1653 | | **[0.1195, 0.2138]** | **[0.1038, 0.2260]** | **[0.0922, 0.2371]** |
| Proportion mediated | 36.6% | | | | |
| ***Pollination*** | | | | | |
| Abundance → Pollination | 0.1176 | 0.0443 | **[0.0588, 0.1739]** | **[0.0440, 0.1938]** | **[0.0300, 0.2110]** |
| Abundance → Richness | 0.5157 | 0.0386 | **[0.4631, 0.5629]** | **[0.4510, 0.5791]** | **[0.4357, 0.5901]** |
| Richness → Pollination | 0.1055 | 0.0402 | **[0.0511, 0.1565]** | **[0.0355, 0.1708]** | **[0.0220, 0.1851]** |
| *Causal mediation analysis* | | | | | |
| Direct effect | 0.1176 | | **[0.0588, 0.1739]** | **[0.0440, 0.1938]** | **[0.0300, 0.2110]** |
| Indirect effect | 0.0540 | | **[0.0259, 0.0810]** | **[0.0173, 0.0884]** | **[0.0105, 0.0963]** |
| Total effect | 0.1710 | | **[0.1205, 0.2223]** | **[0.1005, 0.2329]** | **[0.0944, 0.2522]** |
| Proportion mediated | 31.6% | | | | |

**Table S9.**

**Results of pairwise comparison of richness-ecosystem service relationships according to the methods used to sample pollinators and natural enemies.** A Bayesian hypothesis testing was used to assess the relative statistical evidence in favor of the null hypothesis versus the alternative hypothesis.

| Hypothesis | Estimate difference | Estimate Error | CI lower | CI upper | Evidence Ratio |
|---|---|---|---|---|---|
| **Pollination** | | | | | |
| Active > Passive | -0.02 | 0.10 | -0.18 | Inf | 0.78 |
| **Pest control** | | | | | |
| Active > Passive | 0.04 | 0.08 | -0.01 | Inf | 2.17 |

**Table S10.**

**Results of pairwise comparison of richness-ecosystem service relationships according to the methods used to quantify pollination and pest control services.** A Bayesian hypothesis testing was used to assess the relative statistical evidence in favor of the null hypothesis versus the alternative hypothesis.

| **Hypothesis** | **Estimate difference** | **Estimate Error** | **CI lower** | **CI upper** |
|---|---|---|---|---|
| **Pollination** | | | | |
| Fruit set = Δ Fruit weight | 0.11 | 0.13 | -0.15 | 0.37 |
| Fruit set = Seed set | -0.10 | 0.08 | -0.26 | 0.06 |
| Δ Fruit weight = Seed set | 0.22 | 0.15 | -0.07 | 0.50 |
| **Pest control** | | | | |
| Cage (damage) = Cage (pest abundance) | 0.09 | 0.18 | -0.26 | 0.44 |
| Cage (damage) = Pest damage | 0.17 | 0.15 | -0.12 | 0.47 |
| Cage (damage) = Sentinel experiments | 0.05 | 0.15 | -0.24 | 0.34 |
| Cage (pest abundance) = Pest damage | 0.08 | 0.14 | -0.19 | 0.36 |
| Cage (pest abundance) = Sentinel experiments | -0.04 | 0.13 | -0.30 | 0.22 |
| Pest damage = Sentinel experiments | -0.12 | 0.10 | -0.32 | 0.07 |

**Supplementary Text**

Detailed Acknowledgements

M.D., E.A.M and I.S.D. were supported by EU-FP7 LIBERATION (311781).

M.D., E.A.M, A.H., J.S and I.S.D were supported by Biodiversa-FACCE ECODEAL (PCIN-2014–048).

C.M.K. was supported by an USDA-NIFA Organic Agriculture Research & Extension Award (2015-51300-24155).

D.S.K. was supported by the "National Socio-Environmental Synthesis Center (SESYNC)—National Science Foundation Award DBI-1052875".

D.A.S. was supported by the Environmental Protection Agency, Ireland.

D.K.L. was supported by the United States Dept. Agriculture-NRI grant 2005-55302-16345.

D.A.L. was supported by the US DOE Office of Science (DE-FCO2-07ER64494) and Office of Energy Efficiency and Renewable Energy (DE-ACO5-76RL01830) to the DOE Great Lakes Bioenergy Research Center, and by the NSF Long-term Ecological Research Program (DEB 1027253) at the Kellogg Biological Station and by Michigan State University AgBioResearch.

H.G.S. was supported by The Swedish Research Council Formas & the Strategic Research Area BECC.

J.G. was supported by the Swiss National Science Foundation, the Mercator Fuondation, and ETH Zurich grants. L.M. was supported by EU-FP7 LIBERATION (311781).

H.G. was supported by an USDA Northeast Sustainable Agriculture Research and Extension Award #GNE12-037.

M.J. was supported by the Centre for Biological Control, SLU.

T.T. was supported by the DFG FOR 2432 (India), the project "Diversity Turn" (VW Foundation), the DFG SFB 990 EFForTS and the GIZ-Bioversity project on Peruvian cocoa.

I.B. was supported by Biodiversa-FACCE ECODEAL (PCIN-2014–048).

S.B. acknowledges her Ph.D scholarship by the Deutsche Bundesstiftung Umwelt (German Federal Environmental Foundation).

A.D.M.B. thanks for a Ph.D scholarship financed by The Coordenação de Aperfeiçoamento de Pessoal de Nível Superior - Brasil (CAPES) - Finance Code 001. P.C. was supported by Programa Nacional Apicola (PROAPI).

A.C. research conducted within the Research Unit FOR1246 (KILI project) funded by the German Research Foundation (DFG).

F.D.S.S. was supported by the “Fundação de Apoio à Pesquisa do Distrito Federal”/FAPDF, Brazil (Foundation for Research Support of the Distrito Federal), nº 9852.56.31658.07042016. This study was also financed in part by the Coordenação de Aperfeiçoamento de Pessoal de Nível Superior - Brasil (CAPES) - Finance Code 001.

G.A.G. was funded by the Dutch Ministry of Agriculture, Nature and Food Quality (BO-11-0.11.01-0.51).

J.E. was supported by The Swedish Research Council Formas & the Strategic Research Area BECC.

T.F. was jointly funded by the Netherlands Organization for Scientific Research (research programme NWO-Green) and BASF - Vegetable Seeds under project number 870.15.030. P.F. and C. L. were supported by ANR grant Peerless “Predictive Ecological Engineering for Landscape Ecosystem Services and Sustainability” (ANR-12-AGRO-0006).

B.M.F thanks the Project "Conservation and Management of Pollinators for Sustainable Agriculture, through an Ecosystem Approach", which is supported by the Global Environmental Facility Bank (GEF), coordinated by the Food andAgriculture Organization of the United Nations (FAO) with implementation support from the United Nations Environment Programme (UNEP) and supported in Brazil by the Ministry of Environment (MMA) and Brazilian Biodiversity Fund (Funbio). Also to the National Council for Scientific and Technological Development - CNPq, Brasília-Brazil for financial support to the Brazilian Network of Cashew Pollinators (project # 556042/2009-3) and a Productivity Research Grant (#302934/2010-3).

M.P.D.G. was funded jointly by a grant from BBSRC, Defra, NERC, the Scottish Government and the Wellcome Trust, under the UK Insect Pollinators Initiative.

C.G. was funded in part by the DOE Great Lakes Bioenergy Research Center (DOE Office of Science BER DE-FC02-07ER64494) and by a USDA Agriculture and Food Research Initiative Competitive Grant 2011-67009-30022.

J.H. was supported by CNPq – INCT-IN-TREE (465767/2014-1), CNPq-PVE (407152/2013-0) and Capes for Ph.D scholarship.

A.L.I. was supported by the University of Michigan - Department of Ecology and Evolutionary Biology and the Rackham Graduate School, University of Michigan.

B.K.K. was supported by the German Research Foundation (DFG).

A.M.K. data are part of the EcoFruit project funded through the 2013-2014 BiodivERsA/FACCE-JPI joint call (agreement# BiodivERsA-FACCE2014-74), with the national funder of the German Federal Ministry of Education and Research (PT-DLR/BMBF) (grant# 01LC1403).

T.K. was supported by the Israeli Ministry of Agriculture and Rural Development (grant number 131-1793-14).

T.N.K. was funded in part by the DOE Great Lakes Bioenergy Research Center (DOE Office of Science BER DE-FC02-07ER64494) and by a USDA Agriculture and Food Research Initiative Competitive Grant 2011-67009-30022.

B.M. was partly funded by a scholarship of the German National Academic Foundation (Studienstiftung des deutschen Volkes) and benefited from financial support from the German Science Foundation (DFG Grant CL-474/1-1, ELUC).

R.E.M. was funded by a Specialty Crop Block Grant from the Wisconsin Department of Agriculture, Trade, and Consumer Protection.

M.G.E.M. was supported by an NSERC PGS-D Scholarship and an NSERC Strategic Projects Grant.

S.G.P. was funded jointly by a grant from BBSRC, Defra, NERC, the Scottish Government and the Wellcome Trust, under the UK Insect Pollinators Initiative.

J.A.R. was supported by an USDA grant CA-D-ENM-5671-RR. A.M.S. was supported by The University of Idaho, the USFWS (NMBCA grant F11AP01025), the Perennial Crop Platform (PCP) and CGIAR - Water Land and Ecosystems (WLE).

M.S. was supported by DFG grant FOR 1246 "Kilimanjaro ecosystems under global change: Linking biodiversity, biotic interactions and biogeochemical ecosystem processes".

A.R.S. research was funded by the CS Fund and the United States Army Research Office (ARO).

M.T. was supported by JSPS KAKENHI Grant Number 16H05061.

H.T. was supported by the Environment Research and Technology Development Fund (S-15-2) of the Ministryof the Environment, and JSPS KAKENHI Grant Number 18H02220.

B.F.V. was supported by CNPq – INCT-IN-TREE (#465767/2014-1), CNPq-PVE (407152/2013-0) and and a Productivity Research Grant CNPq (#305470/2013-2).

C.W. was supported by the Deutsche Forschungsgemeinschaft (DFG) (Projekt number 405945293).

B.K.W. was supported by a PhD scholarship from the University of New England and funded by RnD4Profit-14-01-008 ''Multi-scale monitoring tools for managing Australian Tree Crops: Industry meets innovation''.

C.Z.T. was supported by a Severo-Ochoa predoctoral fellowship (SVP-2014-068580) and by the Biodiversa-FACCE project ECODEAL.

W.Z. was supported by CGIAR research program on Water, land and ecosystems (WLE).

Y.Z. was funded by the Division for Earth and Life Sciences of the Netherlands Organization for Scientific Research (grant 833.13.004).
